# Tbr2-Dependent Parallel Pathways Regulate the Development of Distinct ipRGC Subtypes

**DOI:** 10.1101/2025.04.29.651262

**Authors:** Takae Kiyama, Ching-Kang Chen, Halit Y. Altay, Yu-Jiun Chen, Leviette Sigala, Dan Su, Steven Eliason, Brad A. Amendt, Chai-An Mao

## Abstract

The intrinsically photosensitive retinal ganglion cells (ipRGC) are the conduit between the retinas and brain regions responsible for non-image-forming and image-forming vision. In mice, six ipRGC subtypes have been discovered based on morphological characteristics, functions, and molecular profiles. All ipRGCs arise from Tbr2-expressing RGCs during developmental stages and subsequently diverge and differentiate into the six mature, distinct subtypes in adult retinas. However, the cellular and molecular mechanisms controlling the formation and maturation of the six ipRGC subtypes remain elusive. Here, we demonstrate that two *Tbr2*-dependent transcription factors, *Iroquois-related homeobox 1* (*Irx1*) and *T-box containing factor 20* (*Tbx20*), are key downstream transcription factors guiding lineage segregations of Tbr2-expressing RGC into distinct adult ipRGC subtypes. Both factors also control *Opn4* expression. *Irx1* is expressed in the M3, M4, and M5 subtypes, while *Tbx20* is predominantly expressed in M1, M2, M6, and subgroups of M3 and M5. When *Irx1* is ablated during retinal development, *Opn4* expression is significantly reduced in the M3, M4, and M5 ipRGC groups; however, the formation of *Irx1*-expressing ipRGCs is not affected. In contrast, when *Tbx20* is deleted, a significant number of *Tbx20*-expressing cells fail to develop while *Opn4* expression is down-regulated. These findings reveal two parallel transcription cascades downstream of Tbr2 for controlling ipRGC subtype formation, fate divergence, and maintenance in the adult retina.

## INTRODUCTION

In vertebrate retinas, retinal ganglion cells (RGCs) are the first neuronal lineage specified from *Atoh7*-expressing retinal precursor cells^1–5^. Once specified, these newly formed nascent RGCs migrate toward the vitreous side of the retina and further differentiate into more than 40 distinct subtypes, including six types of intrinsically photosensitive RGCs (ipRGCs), each with unique functions, dendritic morphologies and stratification patterns, presynaptic partners, and postsynaptic central projection targets in the brain for different physiological tasks^6–27^. These concerted processes take place around embryonic day 12 (E12) and involve a series of cellular remodeling events and intense cell-cell and cell-environment interactions, which are governed by hierarchical transcriptional regulatory programs and extrinsic factors^28–30^. A significant amount of knowledge has been built regarding the developmental mechanisms regulating RGC specification and differentiation during the early phase of RGC development; however, little is known about the molecular mechanisms that control how nascent RGCs differentiate into the diverse, mature RGC subtypes in the adult retina.

Previously, we discovered T-box transcription factor *T-brain 2* (*Tbr2* or *Eomes*) as the key regulator for the formation and maintenance of ipRGCs^19, 31, 32^. In a follow-up study, we further discovered that all six ipRGC subtypes arise from *Tbr2*-expressing RGCs during development, establishing that Tbr2-expressing RGCs are ipRGCs and that Tbr2 is the key regulator positioned at the very top of the genetic hierarchy governing ipRGC development^22^. However, the molecular mechanisms controlling the lineage segregation of Tbr2-expressing RGCs into six adult ipRGC subtypes remain unknown. To address this question, we focus on exploring the roles of Tbr2-regulated downstream transcription factors in the development of ipRGC subtypes. Using bulk RNA-sequencing (RNA-seq) on E15.5 *WT* and *Tbr2*-mutant retinas, we discovered a small subset of Tbr2-dependent downstream genes, including two transcription regulators: *Irx1* and *Tbx20*. To track the spatiotemporal expression pattern of *Irx1* and *Tbx20* during retinogenesis, we generated two novel knock-in mouse lines, *Irx1^HA3-P2ACreERT2^* and *Tbx20^HA3-P2ACreERT2^*, respectively, and paired them with *Rosa^iAP^* and *Ai9* reporter lines to visualize the identity and to study lineage segregation mechanism of *Irx1*- and *Tbx20*-expressing ipRGC subtypes during development. Furthermore, we also generated several novel knock-in and conditional knockout (KO) mouse lines, including *Irx1^flox^*, *Irx1^LacZ^*, *Tbx20^flox^*, and *Tbx20^LacZ^* alleles, to help unveil their involvement in ipRGC development.

Here, we demonstrate that *Irx1* and *Tbx20* are expressed in distinct subsets of RGCs from embryonic stages onwards and continue to be expressed in distinct types of ipRGCs in adult retina. We show that the onset of Irx1 and Tbx20 marks the beginning of lineage segregation of these ipRGCs in developing retinas. Loss-of-function studies show that *Irx1* is essential for the expression of *Opn4* in M3, M4, and M5 types of ipRGCs, and *Tbx20* is required for the expression of *Opn4* in M1, M2, and M6 types of ipRGCs and is also involved in the formation and maintenance of some of these three ipRGC subtypes. This study reveals two parallel transcriptional pathways downstream of Tbr2 that guide newly developed neurons toward distinct fates, each characterized by unique molecular, morphological, and functional identities.

## RESULTS

### Bulk RNA-seq identified Irx1 and Tbx20 as the candidate Tbr2-dependent downstream transcription factors for ipRGC development

To explore *Tbr2* downstream effector genes that determine ipRGC subtype specification, we conducted bulk RNA-sequencing analysis on embryonic day 15.5 (E15.5) *wild-type* (*Ctrl*) and *Six3-Cre;Tbr2^fx/fx^* (*Tbr2*-*CKO*) retinas to identify differentially expressed genes whose expression was affected in *Tbr2*-*CKO* retinas. By comparing the transcriptomic profiles between these two groups, we uncovered 19 *Tbr2*-dependent genes whose expressions were either down-regulated (13/19) or up-regulated (6/19) in *Tbr2*-*CKO* retinas (Figure 1A). Quantitative analysis using RT-qPCR validated the expression levels of 12 of the down- and 2 of the up-regulated genes in *Tbr2*-*CKO* retinas, which revealed no sex-based differences in relative levels of these genes (Figure 1B). Next, we then reanalyzed the E16 RGC-enriched single-cell RNA-sequencing (scRNA-seq) data from Shekhar et al., 2022 (GEO: GSE185671) and found that *Tbr2*/*Eomes* expression is highly enriched in cluster 3 (C3) and C4, and is detectable at lower levels in C11 and C12 (Extended Figure 1A)^33^. As expected, we found that most of the affected genes in our RNA-seq data are enriched in these clusters except for *Rgs4* (Extended Figure 1A), and that the expression of the up-regulated genes *Lmo2* is distributed across many clusters, and that *Kcnh1* is relatively low in all clusters (Extended Figure 1A). The trend of their correlated expression persists into P0 scRNA-seq data (Geo: GSE185671; Extended Figure 1B), corroborating the use of bulk RNA-seq in uncovering *Tbr2*-dependent genes in embryonic retinas.

**Figure 1.**
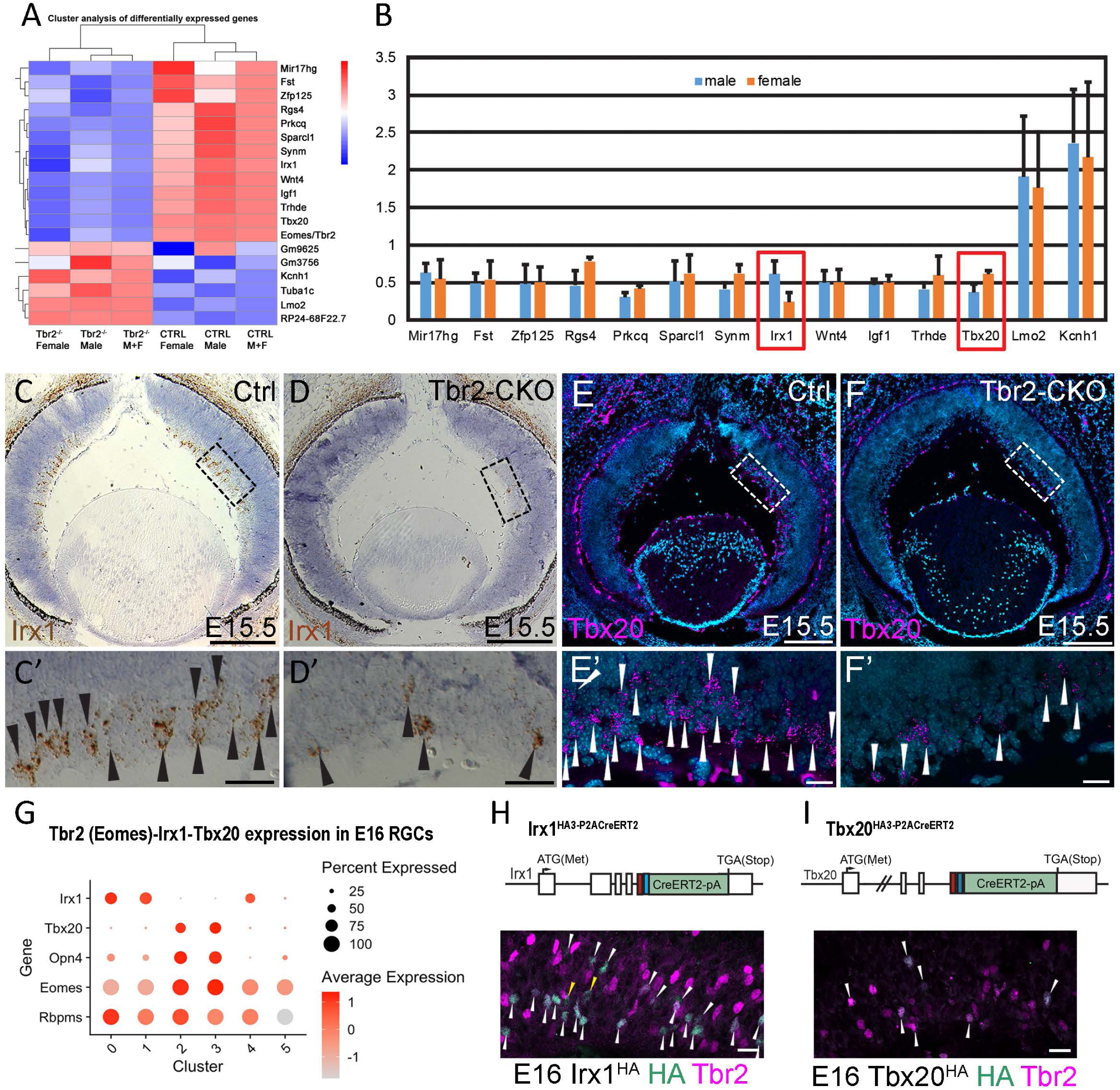
Bulk RNA-seq identified regulatory genes downstream of Tbr2 in developing mouse retinas. **(A)** Heatmap of genes with altered expression levels in the *Tbr2-CKO* retina. **(B)** Validation of RNA-seq data using RT-qPCR between E15.5 *wildtype* and *Tbr2-CKO* retinas. (**C-F**) In situ hybridization of *Irx1*(C, D) and *Tbx20* (E, F) on E15.5 *wildtype* (C, D) and *Tbr2-CKO* (E, F) retinal sections. (C’-F’) Higher magnification images of areas in dashed boxes. Irx1+ and Tbx20+ cells are indicated with arrowheads. **(G)** Reanalysis of single cell-RNA sequencing data for E16 RGCs (Geo dataset: GSE185671). **(H, I)** *Irx1^HA3-P2ACreERT2^* (H) and *Tbx20^HA3-P2ACreERT2^* (I) alleles and retinal sections (E15.5) co-labeled with anti-HA and anti-Tbr2 antibodies. Scale bar: 200 μm in C-F, 50 μm in C’-F’, 20 μm in H and I.

Among these affected genes, we focused on *Irx1* and *Tbx20* due to their broader roles as transcription regulators in several developmental pathways and their closely correlated expression in *Eomes+* clusters in published scRNA-seq data. By in situ hybridization, we were able to establish that *Irx1* and *Tbx20* are both expressed already in the ganglion cells at E15.5 *Ctrl* retinas (Figure 1C, 1C’, 1E and 1E’) and that their expression level was down-regulated in *Tbr2-CKO* retinas (Figure 1D, 1D’, 1F and 1F’), confirming that *Irx1* and *Tbx20* were *Tbr2*-dependent downstream genes. We then subsetted the E16 and P0 scRNA-seq data using a cutoff of Rbpms>1 and Eomes>1, and compared their expression profiles with *Opn4* and *Eomes*. Intriguingly, *Irx1* and *Tbx20* are found in distinct clusters in both datasets (Figure 1G and Extended Figure 1C). In addition, *Pou4f1* expression is relatively weak (if not absent) in those clusters with higher expression levels of *Eomes* and *Opn4* (Extended Figure 1D, 1E). The segregation of *Pou4f1+* and *Eomes+* clusters during RGC development corroborated previous findings that their expressions are mutually exclusive in adult retinas^22^, suggesting that fate segregation of ipRGCs and non-ipRGCs is completed as early as P0. Together, these data raised the possibility that more than one transcription cascade, namely, *Tbr2-Irx1 and Tbr2-Tbx20,* might be involved in ipRGC subtype formation, lineage segregation, and/or regulation of gene expression therein.

### Spatiotemporal expression profile of Irx1 and Tbx20 during retinal development

To track the spatiotemporal expression profiles of *Irx1*, we used CRISPR/Cas9 knockin (KI) technique to generate an *Irx1^HA3-P2ACreERT2^* mouse line in which an *HA3-P2A-CreERT2* reporter cassette was inserted in-frame to *Irx1’s* c-terminal codon (Extended Figure 2A). This new *Irx1* reporting line, labeled as *Irx1^HA^* or *Irx1^CreERT2^* interchangeably in this text according to its usage relevancy, harbors 3 copies of HA tag sequence C-terminally fused with Irx1 protein followed by a releasable CreERT2 produced by self-cleaving 2A peptide through ribosomal skipping (Extended Figure 2A). In parallel, we generated a *Tbx20^HA3-P2ACreERT2^* knock-in mouse line with a similar strategy to monitor *Tbx20*-expressing cells (labeled as *Tbx20^HA^* or *Tbx20^CreERT2^* according to usage, Extended Figure 2B). By co-immunostaining with anti-HA and anti-Tbr2 antibodies on E16.5 *Irx1^HA/+^* and *Tbx20^HA/+^* retinas, we found overlapping HA and Tbr2 signals (Figure 1H and 1I) despite a few non-overlapped ones in *Irx1^HA/+^* sections (yellow arrowheads in Figure 1H).

### Characterization of Irx1-expressing RGCs

Approximately 2500 evenly distributed HA-labeled signals were detected in adult *Irx1^HA/+^* retinas without a skewed distribution in retina quadrants (Figure 2A, 2B). Most of the HA+ signals were also co-labeled with Tbr2 and the pan-RGC marker Rbpms (white arrowheads in Figure 2C), confirming that Irx1 expression was mostly restricted to Tbr2+ RGCs. However, a few weak HA+ cells were negative for Tbr2 (yellow arrowheads in Figure 2C). Because Tbr2-expressing RGCs are essentially ipRGCs, we examined the expression of Opn4, Spp1, and SMI32, the known markers for specific ipRGC subtypes^34, 35^, with HA in *Irx1^HA/+^* retinas. Interestingly, we found that only a small fraction of HA+ cells expressed melanopsin (Figure 2D), suggesting that the majority of Irx1+ RGCs were not the M1 type. These Irx1+ RGCs included Spp1- SMI32- (M5 ipRGC; ~75%) and Spp1+ SMI32+ (M4 ipRGCs; ~22%) (Figure 2E, 2F). Some Spp1- SMI32+ (likely M3 or M6) and Spp1+ SMI32- (likely M2) RGCs were also observed, although they consisted of a very small fraction (~3%). Additionally, we did not detect a spatially distorted retinal expression pattern in these groups (Figure 2F). These data suggested that the majority of Irx1-RGCs were composed of the M4 and M5 subtypes.

**Figure 2.**
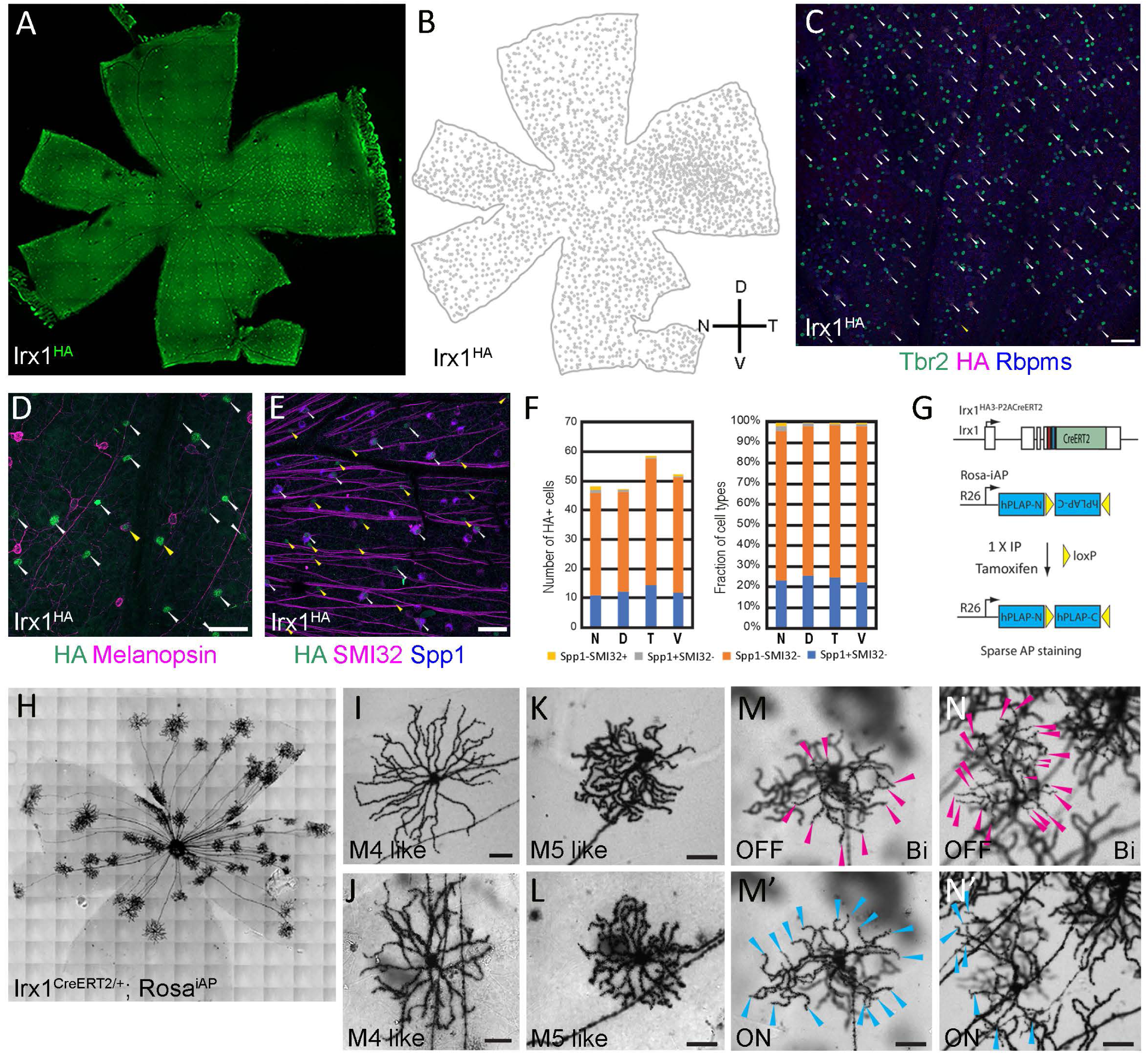
*Irx1*-expressing RGCs. **(A)** Immunofluorescent staining on *Irx1^HA/+^* retinal flatmount with anti-HA antibody. **(B)** Lucida tracing of HA+ cells in A. **(C)** IF staining of *Irx1^HA/+^* retina with anti-HA, anti-Tbr2, and anti-RBPMS antibodies. HA+ Tbr2+ RBPMS+ cells are indicated with white arrowheads. **(D)** IF staining of *Irx1^HA/+^* retina with anti-HA and anti-melanopsin antibodies. HA+ melanopsin+ and HA+ melanopsin-cells are marked with yellow and white arrowheads, respectively. **(E)** IF staining of *Irx1^HA/+^* retina with anti-HA, anti-SMI32, and anti-Spp1 antibodies. SMI32+ Spp1+, SMI32- Spp1-, and SMI32+ Spp1-cells are indicated by white, yellow, and double white arrowheads, respectively. **(F)** The number of HA+ cells expressing Spp1 and/or SMI32 (unit area: 0.33 mm^2^). **(G)** Schematic illustration showing genetically-activated *Rosa^iAP^* reporter by *Irx1^CreERT2^*. **(H-N)** *Irx1*-expressing cells revealed by genetically-directed alkaline phosphatase (AP) labeling. (H) AP staining pattern on flat-mounted *Irx1^CreERT2^; Rosa^iAP^* retina. (I-N’) Representative images showing distinct subtypes of ipRGCs. Scale bars: 50 μm. D: dorsal. N: nasal. T: temporal. V: ventral.

Next, to visualize and determine the types of ipRGC expressing Irx1 by dendritic morphology and retinofugal projections, we bred *Irx1^CreERT2^* with the *Rosa^iAP^* reporter line to produce *Irx1^CreERT2/+^; Rosa^iAP/+^* mice and performed genetic sparse alkaline phosphatase (AP) staining on flat-mounted retinas (Figure 2G, Extended Figure 3A). In all retinas examined (n = 11), we identified 3 morphologically distinct RGC types (Figure 2H-2N). The first type occupied 22% of discernible AP+ RGCs (28/127; Figure 2I, 2J; Extended Figure 3D). They had large, symmetrically distributed dendritic arbors that stratify to the ON layer in the inner plexiform layer (IPL). Morphologically, these cells appeared to be the M4 ipRGCs, or ON α-RGCs^36, 37^. The second RGC type, accounting for 36% of AP+ RGCs (46/127), displayed a smaller symmetrical dendritic arbor stratified to the ON layer in the IPL (Figure 2K, 2L; Extended Figure 3B, 3C). These cells appeared to be the M5 ipRGCs. In the third population of AP+ cells, we found many RGCs (49/127; 49%) whose dendrites stratify to both ON and OFF layers, suggesting that these are the M3 ipRGCs (Figure 2M-2N’; Extended Figure 3E, 3E’). In addition to these ipRGC-like cells, we also detected a small population of OFF-layer stratified RGCs, including a J-RGC (1/127) and a type of RGCs with very dense dendritic arbor (3/127) (Extended Figure 3F, 3G), consistent with the observation that some Irx1+ RGCs did not express Tbr2 (Figure 2C).

To validate the ipRGC subtypes revealed in AP staining, we bred *Irx1^CreERT2^* with the *R26R-tdTomato* (*Ai9*) reporter line to produce *Irx1^CreERT2/+^; Ai9* mice (*Irx1-Ai9*; Extended Figure 4A) and conducted dye-filling and patch-clamp recording according to our previous study on Tbr2+ ipRGCs^22^. Robust tdTomato expression in *Irx1-Ai9* retinas was achieved after two consecutive tamoxifen injections (Extended Figure 4B). By immunostaining, these tdTomato+ cells co-labeled with Irx1-HA expression (Extended Figure 4C), and the RGCs expressing Spp1 appeared to be larger RGCs, presumptively the M4 ipRGCs (white arrowheads, Extended Figure 4C, 4D). Patch-clamp recording in conjunction with biocytin filling confirmed the presence of these three ipRGC types (n=103 Figure 3A), with M4 and M5 being the majority (Fig. 3B-3D, > 80%). Interestingly, for the bi-stratified M3 ipRGCs, not all subtypes with characteristic intrinsic membrane property (IMP) profiles described in the *Tbr2^CreERT2/+^; Ai9* (*Tbr2-Ai9*) mouse retinas were found. The M3s (Figure 6h in Chen *et al.,* 2021)^22^ subtype with no apparent hyperpolarization-activated channels was notably missing, while the other two types (M3n and M3r corresponding to Figure 6i and 6j in Chen *et al.,* 2021, respectively)^22^ were identified (Fig. 3E and 3F). The nomenclature of these subtypes follows the categorizing principles for the M1-ipRGCs by IMP profiles by Chen *et al,* 2021.^22^. The “n” and “r” designations are for cells with prominent hyperpolarization-activated channels, with the “n” designation for the lack of rebound spiking and the “r” for rebound spiking following negative current injections. Moreover, we also encountered a novel M5 subtype with an IMP profile, coined M5nt (no tail) (Figure 3C), in addition to the two M5 subtypes reported in the *Tbr2-Ai9* mice: the M5r (Figure 8d in Chen *et al.*, 2021)^22^ and the M5n (Figure 8f in Chen *et al.*, 2021)^22^. This novel M5nt subtype constituted ~63% of all M5 cells encountered in the *Irx1-Ai9* mice, and in the presence of glutamatergic blockade, 90% of them exhibited robust intrinsic photosensitivity (n= 20, Figure 3C). In summary, Irx1-expressing RGCs are predominantly ipRGCs of the M3, M4, and M5 types.

**Figure 3.**
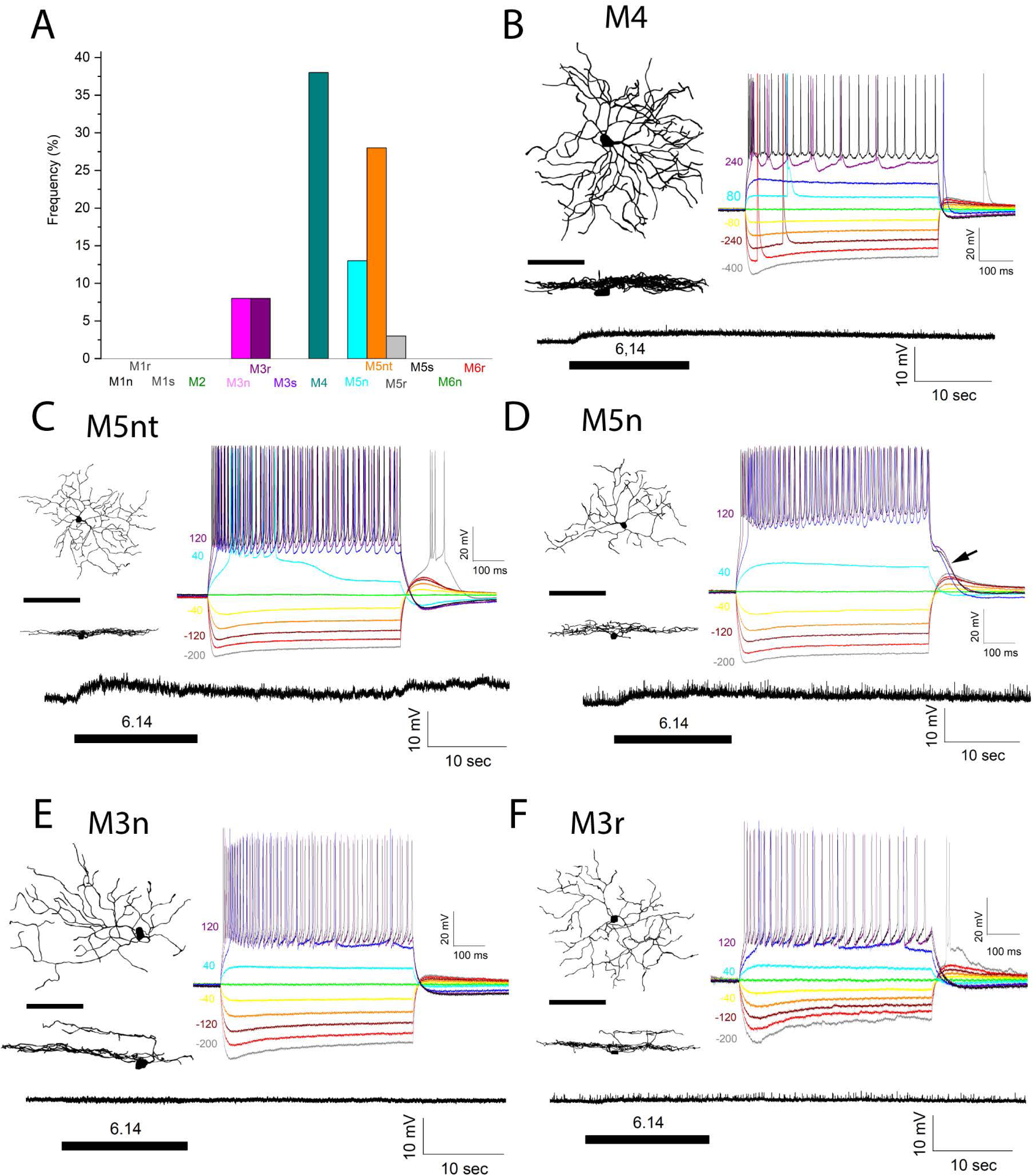
Electrophysiological properties of *Irx1*-expressing ipRGCs in the *Irx1^CreERT2^; Ai9* mouse. **(A)** Frequency of encounter of ipRGC IMP subtypes from 103 randomly targeted tdTomato-positive cells. **(B)** Representative M4 ipRGC with intrinsic membrane property at right, intrinsic photosensitivity at bottom, and virtually reconstructed dendritic morphology at left in the XY (upper) and XZ (lower) planes. **(C)** Representative M5nt ipRGC. **(D)** Representative M5n ipRGC with the characteristic “tail” appearance following the positive current injections indicated by an arrow. **(E)** Representative M3n ipRGC. **(F)** Representative M3r ipRGC. Scale bars for dendritic morphology equal 100 μm. Intrinsic photoresponse was initiated by a 15-sec 470 nm light stimulation (~430 mm in diameter) at 6.14 log R*/rod/sec. Polarities and amounts of current injected to obtain the intrinsic voltage responses are shown at left.

### Characterization of Tbx20-expressing RGCs

One of the most revealing findings in the *Irx1-Ai9* mouse retinas is the absence of M2 and M6 ipRGCs, suggesting that they were specified by other factors. In adult *Tbx20^HA/+^* retinas, approximately 2550 HA-labeled signals were detected across the retina (Figure 4A, 4B). Unlike the case of Irx1, these HA+ signals exhibited a gradient with higher density in the ventral-nasal region and lower density in the dorsal-temporal region (Figure 4A-4C). All HA+ signals were co-labeled with Tbr2 and the pan-RGC marker RBPMS (white arrowheads in Figure 4D), and Tbx20 expression was restricted to Tbr2+ RGCs. Upon examination of the co-expression pattern between melanopsin, Spp1, and HA in *Tbx20^HA/+^* retinas (Figure 4E, 4F), we found that, in the ventral and nasal regions, the majority of Tbx20+ ipRGCs were melanopsin- and Spp1- (Figure 4G). Based on the morphological and recording data described in the next section, we concluded that these are primarily M6 ipRGCs (Figures 4G, 4I and 5B, 5C) characterized by a small but dense dendritic field with numerous ascending and diving dendrites. In contrast, in the dorsal retina, the majority of Tbx20+ ipRGCs were melanopsin+ and Spp1-, suggesting that they might be the M1 type of ipRGCs (Figure 4G). In the temporal retinas, the numbers of these two groups were comparable (Figure 4G). We noted a smaller fraction of melanopsin+ Spp1+ cells in all retinal quadrants, likely to be the M2-like ipRGCs (Figure 5D). These data suggested that Tbx20+ RGCs are consisted of multiple ipRGC subtypes, primarily M1, M2, and M6, in stark contrast to the M3, M4, and M5 ipRGC types found in the *Irx1-Ai9* mice. The type-specific skewed distribution along the ventral-nasal to the dorsal-temporal axis likely reflected the distribution of the abundant M6 ipRGCs labeled in this line.

**Figure 4.**
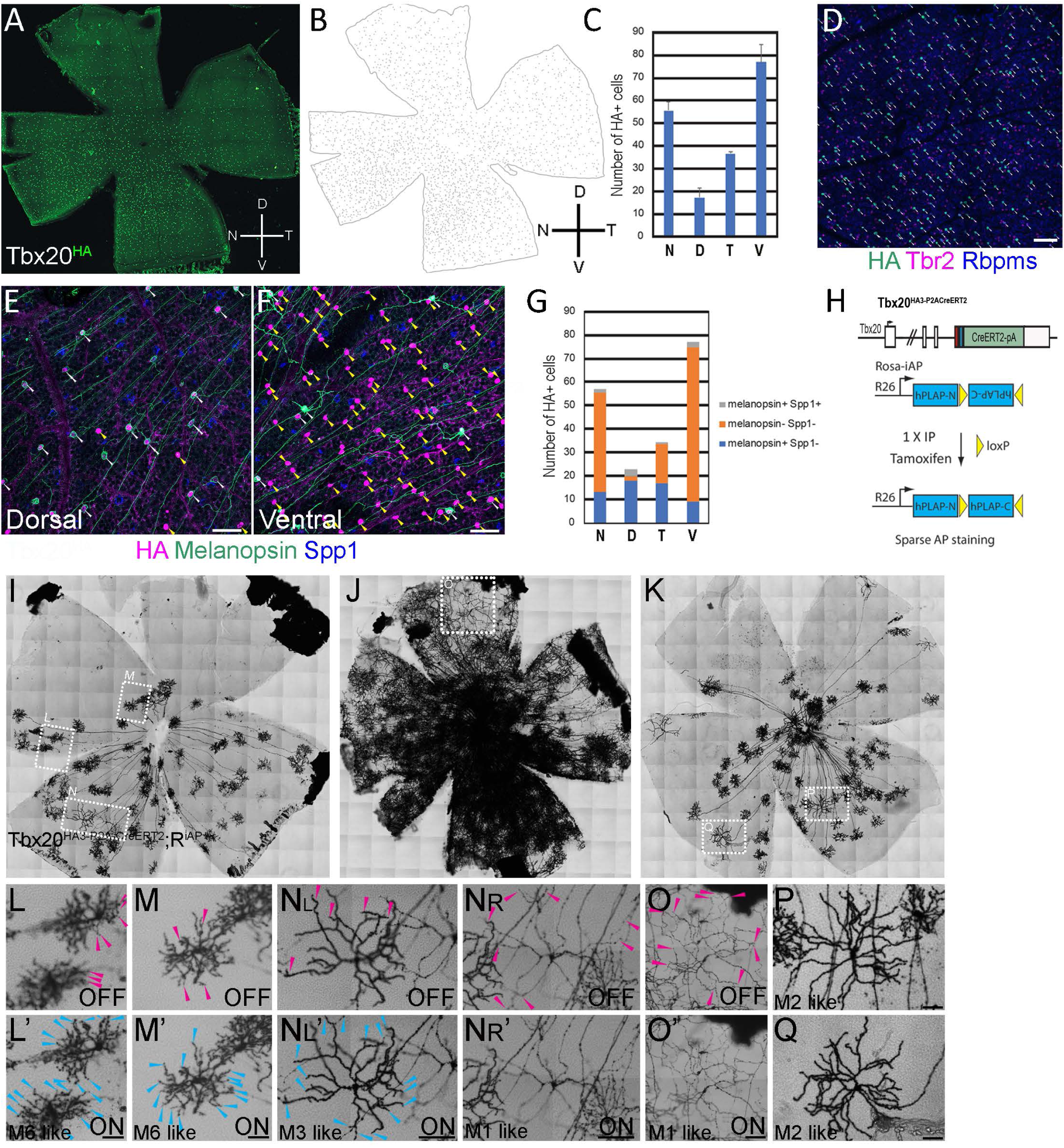
*Tbx20*-expressing ipRGCs. **(A)** IF staining on *Tbx20^HA3/+^* retinal flatmount with anti-HA antibody showing a VN high and DT low expression pattern of Tbx20. **(B)** Lucida tracing of HA+ cells in A. **(C)** Number of HA+ cells in the four quadrants (Unit area = 0.2 mm^2^). **(D)** IF staining of *Tbx20^HA3/+^* retina with anti-HA, anti-Tbr2, and anti-RBPMS antibodies. HA+ Tbr2+ RBPMS+ and HA+ Tbr2-RBPMS+ cells are marked with white and yellow arrowheads, respectively. **(E, F)** IF staining of *Tbx20^HA3/+^* retina in dorsal (E) and ventral (F) regions with anti-HA, melanopsin, and Spp1 antibodies. Melanopsin+ Spp1-, melanopsin-Spp1-, and melanopsin+ Spp1+ cells are indicated by white, yellow, and double white arrowheads, respectively. **(G)** The number of HA+ cells expressing melanopsin and/or Spp1. **(H)** Schematic illustration showing genetically-activated *Rosa^iAP^* reporter by *Tbx20^CreERT2^*. **(I-Q)** Tbx20-expressing cells revealed by genetically directed AP labeling. (I.J, K) AP staining patterns on flat-mounted *Tbx20^CreERT2^; Rosa^iAP^* retina without (I, K) and with (J) 4-OHT exposure. (K-Q) Representative images showing distinct subtypes of ipRGCs in I-K. ON and OFF-layer dendritic terminals are indicated with blue and red arrowheads, respectively. Scale bar: 50 μm in D, E, F, K, L’, M’, P, and Q; 100 μm in N and O’. D: dorsal. N: nasal. T: temporal. V: ventral.

**Figure 5.**
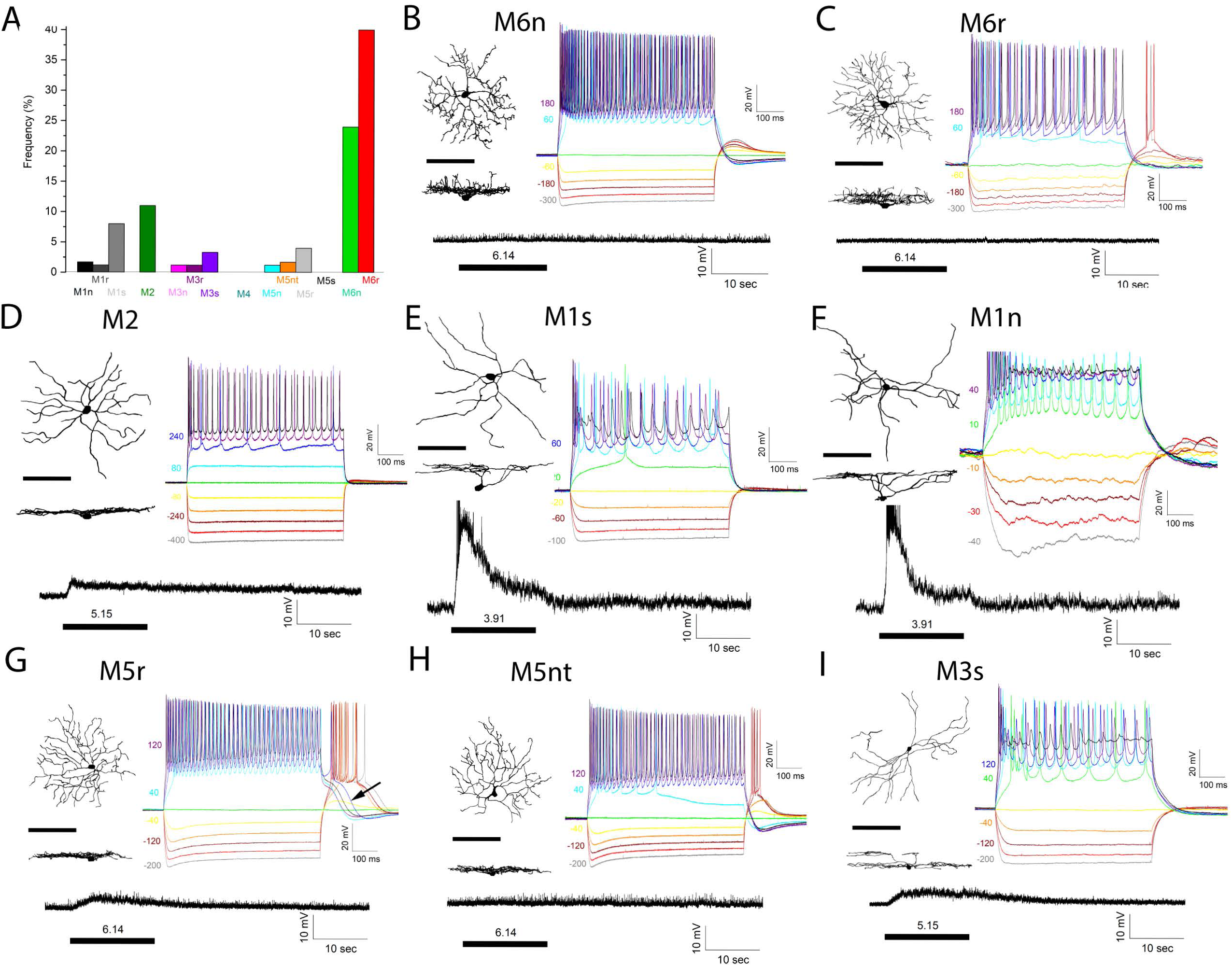
Electrophysiological properties of *Tbx20*-expressing ipRGCs in the *Tbx20^CreERT2^; Ai9* mouse. **(A)** Frequency of encounter of ipRGC IMP subtypes from 181 randomly targeted tdTomato-positive cells. **(B)** Representative M6n ipRGC with intrinsic membrane property at right, intrinsic photosensitivity at bottom, and virtually reconstructed dendritic morphology at left in the XY (upper) and XZ (lower) planes. **(C)** Representative M6r ipRGC. **(D)** Representative M2 ipRGC. **(E)** Representative M1s ipRGC. **(F)** Representative M1n ipRGC. **(G)** Representative M5r ipRGC with the characteristic “tail” appearance following the positive current injections indicated by an arrow. **(H)**. Representative M5nt ipRGC. **(I)**. Representative M3s ipRGC. Scale bars for dendritic morphology equal 100 μm. Intrinsic photoresponse was initiated by a 15-sec 470 nm light stimulation (~430 mm in diameter) at 3.91 to 6.14 log R*/rod/sec as indicated. Polarities and amounts of current injected to obtain the intrinsic voltage responses are shown at left.

To morphologically ascertain the ipRGC types that express Tbx20, we generated *Tbx20^CreERT2/+^; Rosa^iAP/+^* mice and performed sparse AP staining on flat-mounted retinas (Figure 4H, Extended Figure 5A). The *Tbx20^CreERT2/+^; Rosa^iAP/+^* mice exhibited some levels of leaky expression without tamoxifen treatment (Figure 4I, 4K), which became evident around post-natal days 10. In adults, a diluted tamoxifen injection via the IP route (one round of 5 μg/g body weight, 4-fold less than the standard dose), numerous AP+ signals appeared to cover the entire retina, preventing a clear examination of the AP-marked dendrites (Figure 4J, Extended Figure 5B, 5C). Therefore, the AP analysis was primarily conducted on the noninjected adult eyes. In all retinas examined (n = 10), most AP+ RGCs were localized in the ventral-nasal retinas, resembling the spatial distribution pattern of melanopsin-Spp1-cells described above (compare Figure 4I, 4K with 4B, 4G). These RGCs appeared as small cells with dense dendritic arbors ramified to the lower half of IPL (or sublaminar B) with a few fine dendrites extending upward into sublaminar A in the IPL (Figure 4L, 4M). Morphologically, these small, bushy, bi-stratified cells appeared similar to the M6-ipRGCs described previously^38, 39^. Occasionally, the M3 type of bi-stratified cells were observed, as well as the M1 types whose terminal dendrite stratified to the uppermost OFF layer near the boundary between the IPL and inner nuclear layer (INL). These cells can readily be identified in the noninjected retinas, but obscured in the tamoxifen-injected retinas (Figure 4I-4K, 4N-4O’). Additionally, we also detected cells with ON-layer stratified dendrites, but their dendritic arbors appear less dense compared to those of the M4 types found in Irx1+ ipRGCs. These cells likely represented the melanopsin+ Spp1+ M2 type of ipRGCs (Figure 4P, 4Q).

To further characterize the cell types revealed by AP staining, we bred *Tbx20^CreERT2^* into the *Ai9* reporter line to produce the *Tbx20^CreERT2/+^; Ai9* mice (*Tbx20-Ai9*, Extended Figure 6A) for dye-filling and patch-clamp recording. Robust tdTomato expression from *Tbx20^CreERT2/+^; Ai9* retinas was observed with one tamoxifen injection (Extended Figure 6B). By immunostaining, these tdTomato+ cells were co-labeled with the Tbx20-HA signal (Extended Figure 6C). Randomly targeted patch-clamp recordings of 181 tdTomato+ cells revealed 19 M1, 20 M2, 10 M3, 13 M5, 117 M6 ipRGCs (Figure 5A) and two WAC3 (data not shown) previously described in the *Tbr2^CreERT2/+^; Ai9* retinas^22^. Therefore, the majority (86%) of RGC types identified by patch-clamp recordings among *Tbx20-Ai9* cells were M1, M2, and M6 ipRGCs, consistent with the results obtained from AP staining. (Figure 5B-5F). The most abundant cell type was the M6, which we divided into two subtypes, namely, M6n and M6r, based on the absence and presence of rebound spiking in their IMP profiles, respectively (Figure 5B and 5C). Of the three M1 subtypes (M1r, M1n, and M1s) described in *Tbr2-Ai9* retinas^22^, the majority found here in *Tbx20-Ai9* was the M1s subtype (~74%), with a few labeled M1r and M1n cells constituting 11% and 15%, respectively (Figure 5A, 5E, 5F). The appearance of M3 and M5 ipRGCs was unexpected but nonetheless understandable as they might be completely masked by the intense AP signal, particularly on the ventral side (Figure 4J). Intriguingly, the IMP and intrinsic photosensitivity of the Tbx20-expressing M3 and M5 ipRGCs differed from those found in the *Irx1-Ai9* mice. While the M3n and M3r subtypes were found in both lines, those in the *Irx1-Ai9* mice displayed barely detectable intrinsic photosensitivity (Figure 3F). Moreover, the M3s subtype was found only in the *Tbx20-Ai9* retinas, and together with the less abundant M3n and M3r, all were notably intrinsically photosensitive (e.g., Figure 5I). Among the four M5 subtypes, M5r, M5n, and M5nt were found in both lines (compare Figure 3A with Figure 5A), but their relative abundance varies. The M5s subtype (corresponding to Figure 8e in Chen *et al.*, 2021; see discussion) was noticeably missing in both lines. Moreover, the intrinsic photosensitivity of the M5nt subtype found in the *Tbx20-Ai9* mice was barely detectable, while those in the *Irx1-Ai9* mice were (compare Figure 5H with Figure 3C). Irx1 and Tbx20 may therefore differentially modulate the expression of genes involved in the phototransduction pathways underlying intrinsic photosensitivity in specific ipRGC types they specified.

### Axonal projection of Irx1- and Tbx20-expressing RGCs in the brain

To determine the retinofugal projection of Irx1+ and Tbx20+ RGCs, we examined AP staining patterns on brain sections from *Irx1^CreERT2/+^; Rosa^iAP/+^* or *Tbx20^CreERT2/+^; Rosa^iAP/+^* mice to which 4-OHT was injected into the right eyes (Figure 6A, 6E).

**Figure 6.**
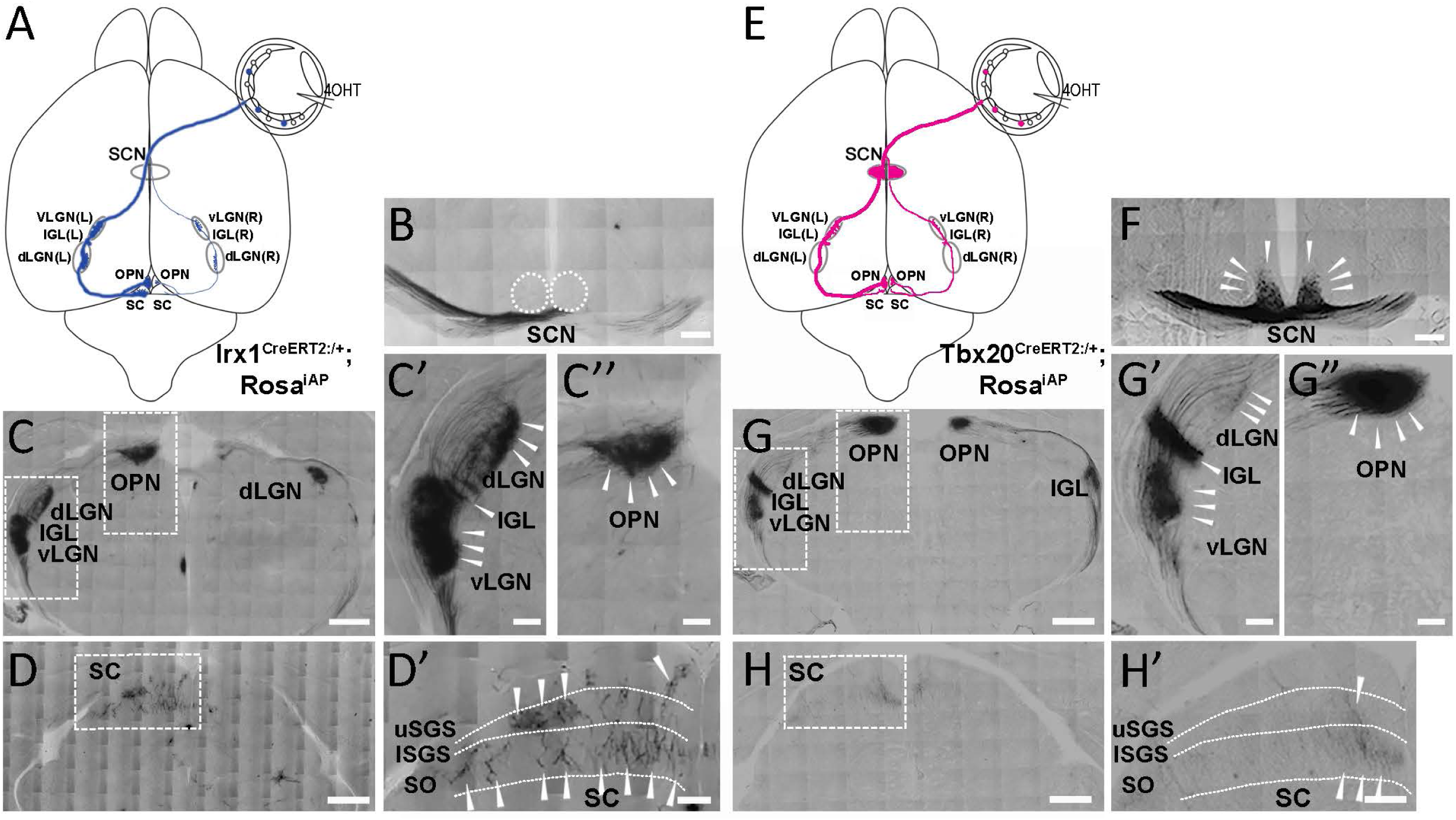
Distinct central projection patterns from Irx1- and Tbx20-expressing ipRGCs in the brain. **(A)** Schematic dorsal view of an *Irx1^CreERT2/+^; Rosa^iAP/+^* mouse brain whose right eye was injected with 4-OHT. The major Irx1+ RGC axonal projections are labeled in blue. **(B)** AP signal was undetectable in SCN. **(C-D”)** Intense AP staining on brain sections showing RGC axon projection in the contralateral side of vLGN, IGL, dLGN, and SC. **(E)** Schematic dorsal view of an *Tbx20^CreERT2/+^; Rosa^iAP/+^* mouse brain whose right eye was injected with 4-OHT. The major Tbx20+ RGC axonal projections are labeled in magenta. **(G-H”)** Intense AP staining on brain sections showing RGC axon projection in SCN, vLGN, IGL, and OPN. Sparse fibers were seen in dLGN and SC. dLGN: dorsal lateral geniculate nuclei; IGL: inter-geniculate leaflet; SC: superior colliculus; vLGN: ventral lateral geniculate nuclei. Scale bars: 500 μm in C, D, G, and H; 100 μm in C’, C’’, G’, G”; 200 μm in B, D’, F, and H”.

For Irx1+ RGCs, AP staining was non-detectable in the suprachiasmatic nuclei (SCN) (Figure 6B), consistent with the finding that Irx1 is not expressed in adult M1 and M2 ipRGCs, whereas intense AP signals were detected in the contralateral side (left) of the ventral lateral geniculate nuclei (vLGN), intergeniculate leaflet (IGL), the inner shell of the dorsal lateral geniculate nuclei (dLGN), and the olivary pretectal nuclei (OPN) (Figure 6C-6C”). In the superior colliculus (SC), AP signals showed a columnar and laminar pattern (Figure 6D, 6D’). Axon arbors projected to different laminae, with relatively weaker signals in the superficial zonal layer (uSGS: upper stratum griseum superficiale) and more intense signals in the inner SGS (ISGS) layer and the deeper optic layer (SO: stratum opticum), spanning across the medial to lateral SC (Figure 6D’)^40^. The axons projecting to the superficial layer likely originated from J-RGCs, while those in the deeper layers likely came from M4 cells^7, 16, 41^.

For Tbx20+ RGCs, sparse AP staining was detected in the SCN (Figure 6F), indicating innervation from the M1 and M2 subtypes of ipRGCs. More intense AP signals were detected in the vLGN, IGL, and OPN (Figure 6G-6G”). In contrast, relatively sparse and weak AP signals were seen in dLGN and SC (Figure 6G’, 6H’). On the ipsilateral side, a similar pattern of AP+ signal was detected, primarily due to the leaky AP expression from mainly the M6 type of ipRGCs of the left retinas.

### Lineage segregation of Irx1- and Tbx20-expressing ipRGCs during development

Based on IF staining, genetic sparse labeling, and recording/dye-filling data, we concluded that Irx1+ RGCs consist mainly of M3, M4, and M5 ipRGCs, while Tbx20+ RGCs are primarily composed of M1s, M2, and M6 with a minor presence of subtypes of M3 and M5 ipRGCs. To test whether Irx1 and Tbx20 might be co-expressed in some RGCs in the adult retina, we generated an *Irx1^HA/+^; Tbx20^LacZ/+^* mouse. It is worth noting that the *Tbx20^LacZ^* allele was derived from the *Tbx20^flox^* allele through germline CMV-Cre-induced recombination (Extended Figure 7A). When *Tbx20^LacZ^* and *Tbx20^HA^* alleles were bred together (*Tbx20^HA/LacZ^*), nearly all HA+ cells were also LacZ+ (>98%; Extended Figure 7B-7D) and hence LacZ expression is a suitable surrogate for Tbx20 expression. IF staining using anti-HA and -LacZ antibodies on P30 *Irx1^HA/+^; Tbx20^LacZ/+^* retinas showed that these two groups of cells were not present in the same cells in the adult retina (Figure 7A, Extended Figure 8). Next, we test whether Irx1 and Tbx20 are co-expressed during embryonic stages using ISH with Irx1 and Tbx20 probes (Figure 7B). We found in E16.5 retinas that *Irx1* and *Tbx20* were detected in distinct RGCs (white dotted circles) with a small fraction of RGCs coexpressing both genes (yellow dotted circles). However, these RGCs consistently exhibited higher expression of one gene over the other, suggesting that they were being segregated into distinct developmental lineages and that their fates were becoming terminally set.

**Figure 7.**
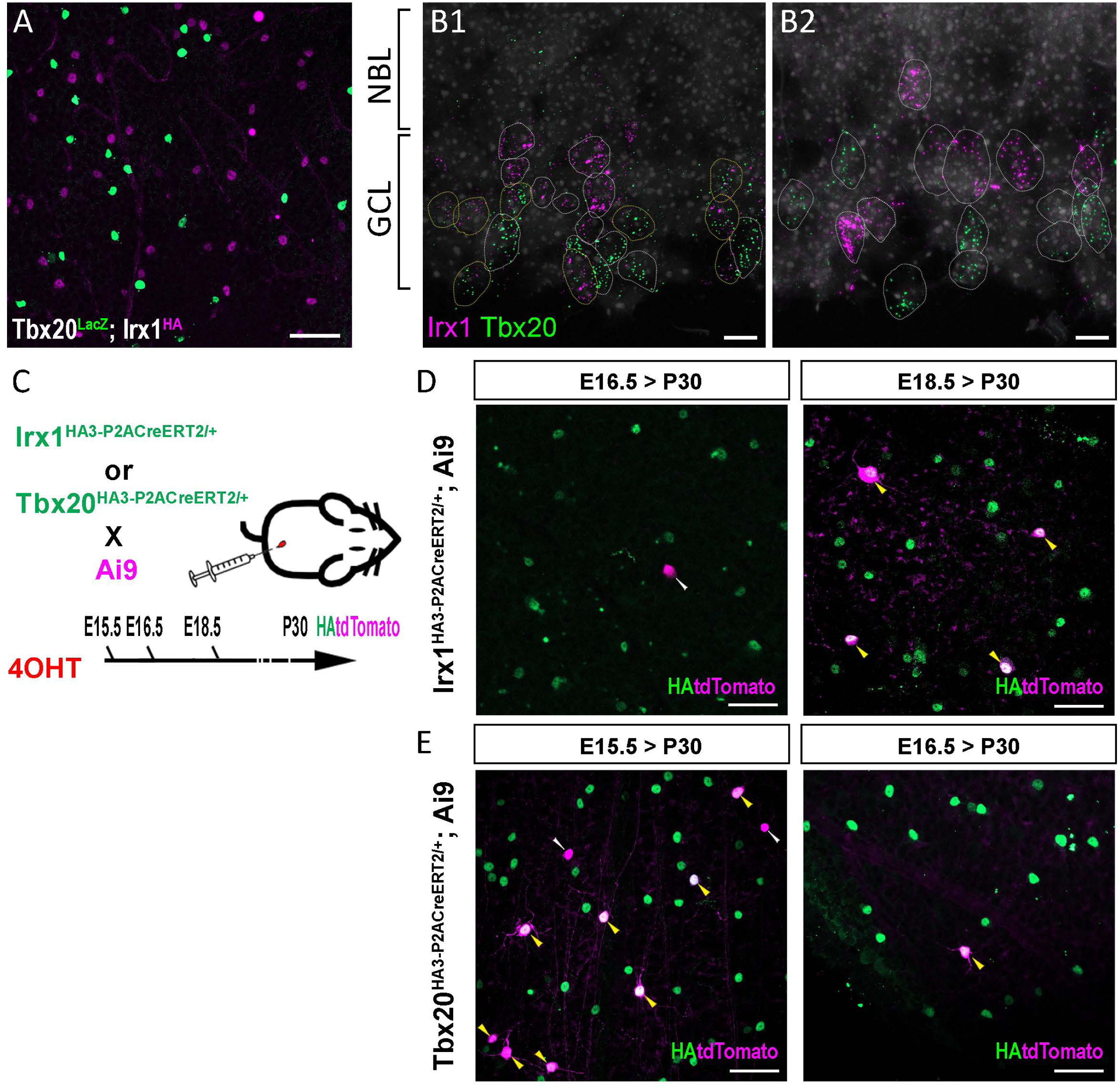
Fate mapping of Irx1- and Tbx20-expressing RGCs. **(A)** IF staining on *Tbx20^LacZ/+^; Irx1^HA/+^* retina using anti-LacZ and HA antibodies showing near complete segregation of Irx1- and Tbx20-expressing RGCs. **(B)** Schematic illustration showing the fate mapping strategy. **(C)** IF staining of HA and tdTomato signals collected from flat-mounted P30 *Irx1^HA3-P2ACreERT2/+^; Ai9* retinas exposed to 4-OHT at E16.5 (left) or E18.5 (right). **(D)** IF staining of HA and tdTomato signals on flat-mounted P30 *Tbx20^HA3-P2ACreERT2/+^; Ai9* retinas exposed to 4-OHT at E15.5 (left) or E16.5 (right). Scale bars: 50 μm.

To determine the developmental stages at which the Irx1-expressing RGCs were fate-determined, we conducted fate mapping for Irx1-expressing cells using the *Irx1*-*Ai9* mice (Figure 7C). We injected 4-OHT into pregnant mice at E18.5 (Figure 7C), and conducted co-IF staining for tdTomato and HA on the P30 retinas from the offspring harboring the *Irx1*-*Ai9* genotype. HA and tdTomato signals completely overlapped, suggesting Irx1-expressing cells in E18.5 retinas were already fate-determined (N = 28; Figure 7D). However, exposure of the *Irx1*-*Ai9* retinas to 4-OHT at E16.5 resulted in ~13% of tdTomato signals lacking HA expression, indicating that Irx1-expressing RGCs (N = 2/15; Figure 7D), including M3, M4, and M5, had acquired their terminal fate between E16.5 and E18.5. Applying the same approach to *Tbx20*-*Ai9* mice, we found that HA+ and tdTomato+ signals completely overlapped in *Tbx20*-*Ai9* retinas exposed to 4-OHT at E16.5 (N = 73; Figure 7E). This overlapped expression pattern was also observed in *Tbx20^CreERT2/+^*; *Ai9* retinas exposed to 4-OHT at E18.5 (data not shown). In contrast, exposure of the *Tbx20*-*Ai9* retinas to 4-OHT at E15.5 resulted in ~20% of tdTomato+ signals not overlapping with HA+ cells (N = 8/39; Figure 7E), indicating that Tbx20-expressing RGCs acquired their terminal fate slightly earlier than Irx1-expressing RGCs, between E15.5 and E16.5.

### Irx1 controls the expression levels of *Opn4* but is dispensable for the formation of Irx1-expressing ipRGCs

To examine the roles of *Irx1* in the expression of *Opn4* and the formation of the ipRGC subset expressing *Irx1*, we first bred *Six3-Cre*, a *Cre* line highly expressed in embryonic retina, with an *Irx1^flox^* allele to generate *Irx1^fx/+^* and *Six3-Cre; Irx1^fx/fx^* littermates for comparison. By IF staining with an anti-melanopsin antibody, we observed a clear reduction in melanopsin signals (Extended Figure 9) in the mutants, suggesting that Irx1 controls *Opn4* expression levels in a subset of ipRGCs. We then bred *Opn4^Cre^*, a *Cre* line activated in newly developed ipRGCs^25^, with *Irx1^fx^* and *Irx1^LacZ^* alleles and tested whether *Opn4* expression was also down-regulated in *Opn4^Cre^; Irx1^LacZ/fx^* versus *Irx1^LacZ/+^* control retinas (Figure 8A). In all three postnatal ages we examined (P7, P21, and P60), a conspicuous and quantitative reduction in melanopsin/*Opn4* expression was seen in all retinal quadrants (Figure 8B, 8C, Extended Figure 10), further ascertaining the essential role of Irx1 for *Opn4* expression. This finding was highly significant because other than *Tbr2*, there has not been a single transcriptional factor shown to regulate *Opn4* expression levels *in vivo*^19, 22, 31^.

**Figure 8.**
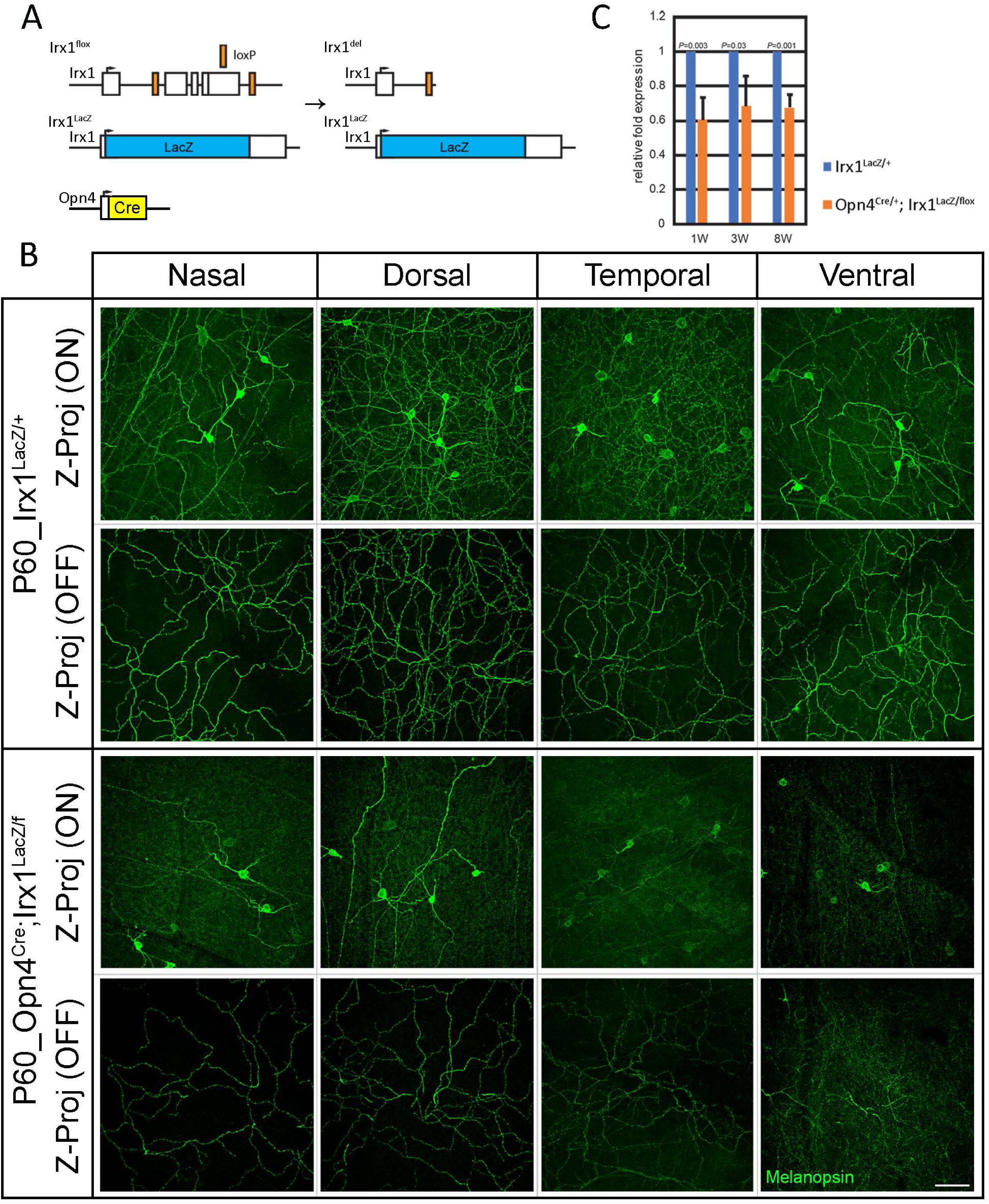
Irx1 controls the expression of *Opn4* but not the formation of Irx1-expressing ipRGCs. **(A)** Schematic diagram showing *Irx1^flox^* and *Irx1^LacZ^* alleles and the conditional *Irx1*-knockout by *Opn4^Cre^*. **(B)** IF staining on *Irx1^LacZ/+^* and *Opn4^Cre^; Irx1^LacZ/fx^* retinas using anti-melanopsin antibody showing a drastic reduction of melanopsin intensity in the ON and OFF sublaminae. The ON-layer images are orthogonal Z-projections of confocal images focused on the ON sublaminae, and the OFF-layer images are orthogonal Z-projections of confocal images focused on the OFF sublaminae. **(C)** Quantitative RT-PCR of *Opn4* in *Irx1^LacZ/+^* and *Opn4^Cre^; Irx1^LacZ/fx^* retinas at 1, 3, and 8 weeks old. Scale bar: 50 μm.

To further distinguish whether the reduction in melanopsin IF signals was due to the down-regulation of *Opn4* expression and/or the loss of *Irx1*-expressing ipRGCs during development, we introduced the *Irx1^LacZ^* allele to generate *Irx1^lacZ/+^* and *Six3-Cre; Irx1^LacZ/fx^* littermates, using LacZ expression as a readout to quantify the number of *Irx1*-expressing RGCs. On X-gal stained flat-mounted retinas, similar numbers of LacZ+ cells were detected in both *Irx1^lacZ/+^* and *Six3-Cre; Irx1^LacZ/fx^* retinas (Extended Figure 11), indicating that *Irx1* is dispensable for the formation of *Irx1*-expressing RGCs but instead required for *Opn4* expression. Together, these data demonstrated that a lineal *Tbr2-Irx1-Opn4* transcriptional cascade controls the expression of *Opn4* in Irx1-expressing ipRGCs.

### Tbx20 controls the expression levels of *Opn4* and the development of the Tbx20-expressing ipRGCs

Until now, the roles of *Tbx20* in the expression of *Opn4* and the development of the ipRGC subset expressing *Tbx20* were obscure, and hence we bred *Six3-Cre* with a *Tbx20^flox^* allele to generate *Six3-Cre; Tbx20^fx/+^* and *Six3-Cre; Tbx20^fx/fx^* littermates for comparison (Figure 9A). By IF staining with an anti-melanopsin antibody and analyzing signal strength akin to what was done for *Irx1*-mutant, we similarly found an obvious reduction in the levels of melanopsin IF signals across all regions (Figure 9B). Quantitative RT-PCR was then performed and confirmed *Opn4* expression was reduced by approximately 25% in the *Tbx20*-mutant (Figure 9C), suggesting that Tbx20, too, controlled *Opn4* expression levels in Tbx20-expressing ipRGCs. To further determine whether this reduction was due to the down-regulation of *Opn4* expression or the loss of Tbx20-expressing RGCs, we compared the LacZ+ cells between *Six3-Cre; Tbx20^fx/+^* and *Six3-Cre; Tbx20^fx/fx^* retinas (Extended Figure 12A, 12B), and found that the number of LacZ+ cells in the mutants decreased to approximately 50% of control (Extended Figure 12C), suggesting that *Tbx20* is essential also for the formation of Tbx20-expressing cells during development. Future work will determine whether the reduction occurs in all major Tbx20+ ipRGCs or is restricted to a specific subset.

**Figure 9.**
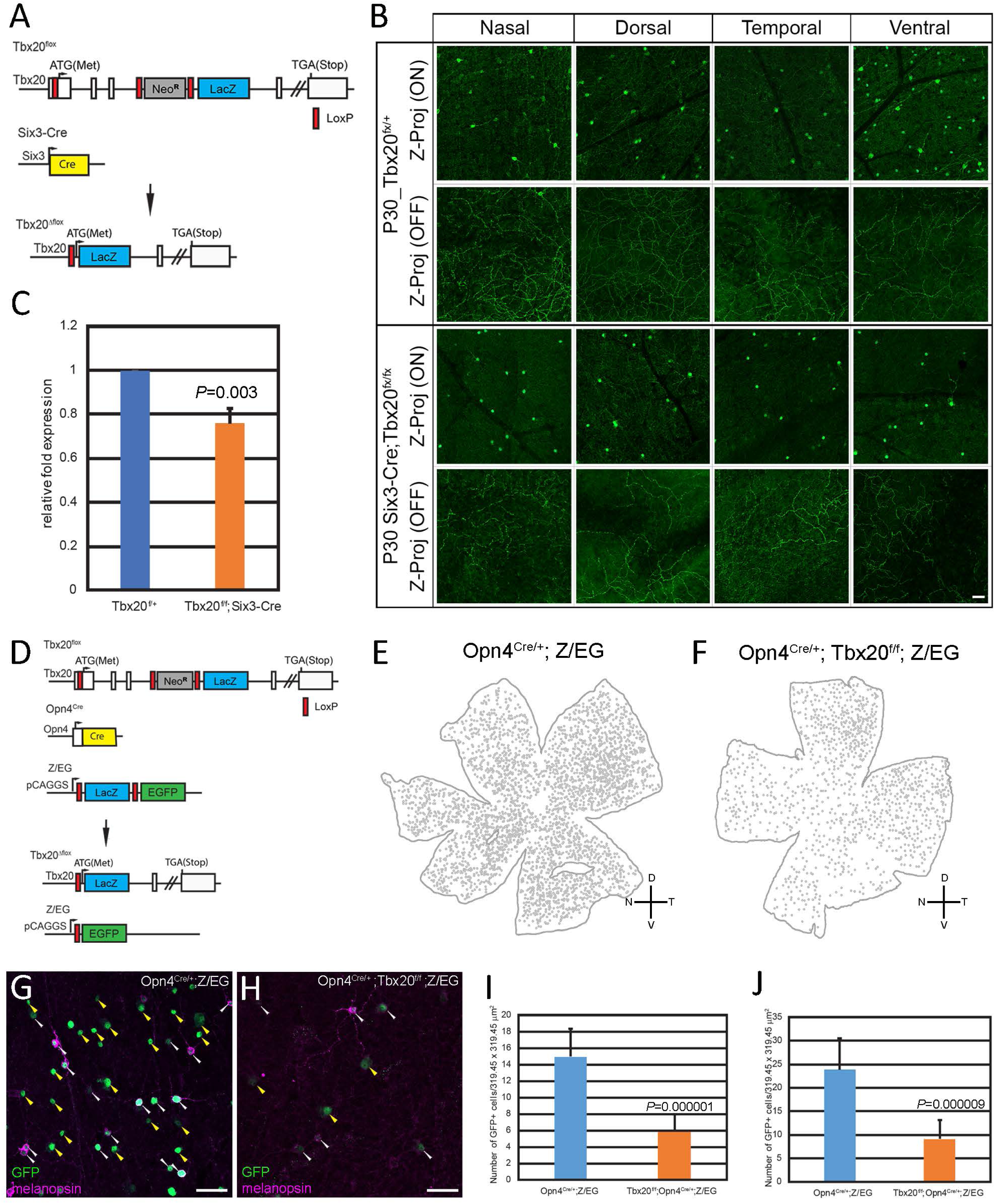
Tbx20 regulates the formation of a subset of ipRGCs and the expression of *Opn4*. **(A)** Schematic illustration showing *Tbx20^flox^* and *Tbx20^LacZ^* (derived from *Tbx20^flox^* upon Cre deletion) alleles and conditional *Tbx20* knockout by *Six3-Cre*. **(B)** Quantitative RT-PCR of *Opn4* in 3-week-old *Tbx20^fx/+^* and *Six3-Cre; Tbx20^fx/fx^* retinas. **(C)** IF staining on *Tbx20^fx/+^* and *Six3-Cre; Tbx20^fx/fx^* retinas with anti-melanopsin antibody showing a drastic reduction of melanopsin intensity in the ON and OFF sublaminae. The ON-layer images are orthogonal Z-projections of confocal images focused on the ON sublaminae, and the OFF-layer images are orthogonal Z-projections of confocal images focused on the OFF sublaminae. **(D)** Schematic illustration showing conditional knockout of *Tbx20* and activation of *Z/EG* reporter by *Opn4^Cre^*. **(E. F)** Lucida tracing of GFP+ cells on *Opn4^Cre/+^; Z/EG* (E) and *Opn4^Cre/+^; Tbx20^fx/fx^; Z/EG* (F) retinas. **(G. H)** IF staining on the ventral regions of *Opn4^Cre/+^; Z/EG* (G) and *Opn4^Cre/+^; Tbx20^fx/fx^; Z/EG* (H) retinas with anti-GFP and anti-melanopsin antibodies. **(I. J)** Number of GFP+ cells in *Opn4^Cre/+^; Z/EG* and *Opn4^Cre/+^; Tbx20^fx/fx^; Z/EG* retina with (I) and without (J) melanopsin signals. Scale bar: 50 μm.

The essential role of *Tbx20* in ipRGC development is reminiscent of the role of Tbr2 and prompted us to examine whether *Tbx20* is also required for the survival of Tbx20-expressing ipRGCs in adults. We bred *Opn4^Cre/+^* with *Tbx20^fx^* and *Z/EG* alleles to generate *Opn4^Cre/+^; Z/EG* control and *Opn4^Cre/+^; Tbx20^f/f^; Z/EG* mutant mice (Figure 9D), and conducted IF staining with an anti-GFP antibody to compare the number of GFP-marked ipRGCs. The number of GFP+ ipRGCs in the mutant was reduced by approximately 66% in the entire retina (compare Figure 9E and 9F). Additionally, the numbers of GFP+ melanopsin+ and GFP+ melanopsin-ipRGCs were reduced (Figure 9G-9J). Together, these data concluded that a hierarchical *Tbr2-Tbx20-Opn4* transcriptional cascade is involved in controlling *Opn4* expression in Tbx20-expressing ipRGCs and also plays an essential role in the formation and maintenance of these ipRGCs.

### Interaction of Irx1 and Tbx20 during ipRGC development

The diminished *Opn4* expression in both *Irx1*- and *Tbx20*-mutant retinas and their transient co-expression in the E16.5 retina prompted us to test whether they interact genetically to control *Opn4* expression and/or the development of ipRGCs. To do this, we bred *Irx1^flox^* and *Tbx20^flox^* alleles with *Six3-Cre* to generate *Irx1^fx/+^; Tbx20^fx/+^* (Ctrl) and *Six3-Cre; Irx1^fx/fx^; Tbx20^fx/fx^* double knockout (dCKO) mice. By IF staining, we found that the number of melanopsin+ ipRGCs was significantly reduced (compared Figure 10A to 10B, Extended Figure 13). Notably, the number of melanopsin+ Spp1+ ipRGCs (M2 and M4; Ctrl: 355.67±28.89 vs dCKO: 64.33±1.52) (Figure 10A1, 10B1) decreased more drastically than that of melanopsin+ Spp1-ipRGCs (mainly M1; Ctrl: 813.33±85.45 vs dCKO: 356.33±69.62) (Figure 10A2, 10B2), suggesting that Irx1 and Tbx20 do play a synergistic role in directing ipRGC development. These data also suggest that there exists additional albeit minor transcriptional pathways downstream of Tbr2 that support ipRGC development in the absence of *Irx1* and *Tbx20* (Figure 10D).

**Figure 10.**
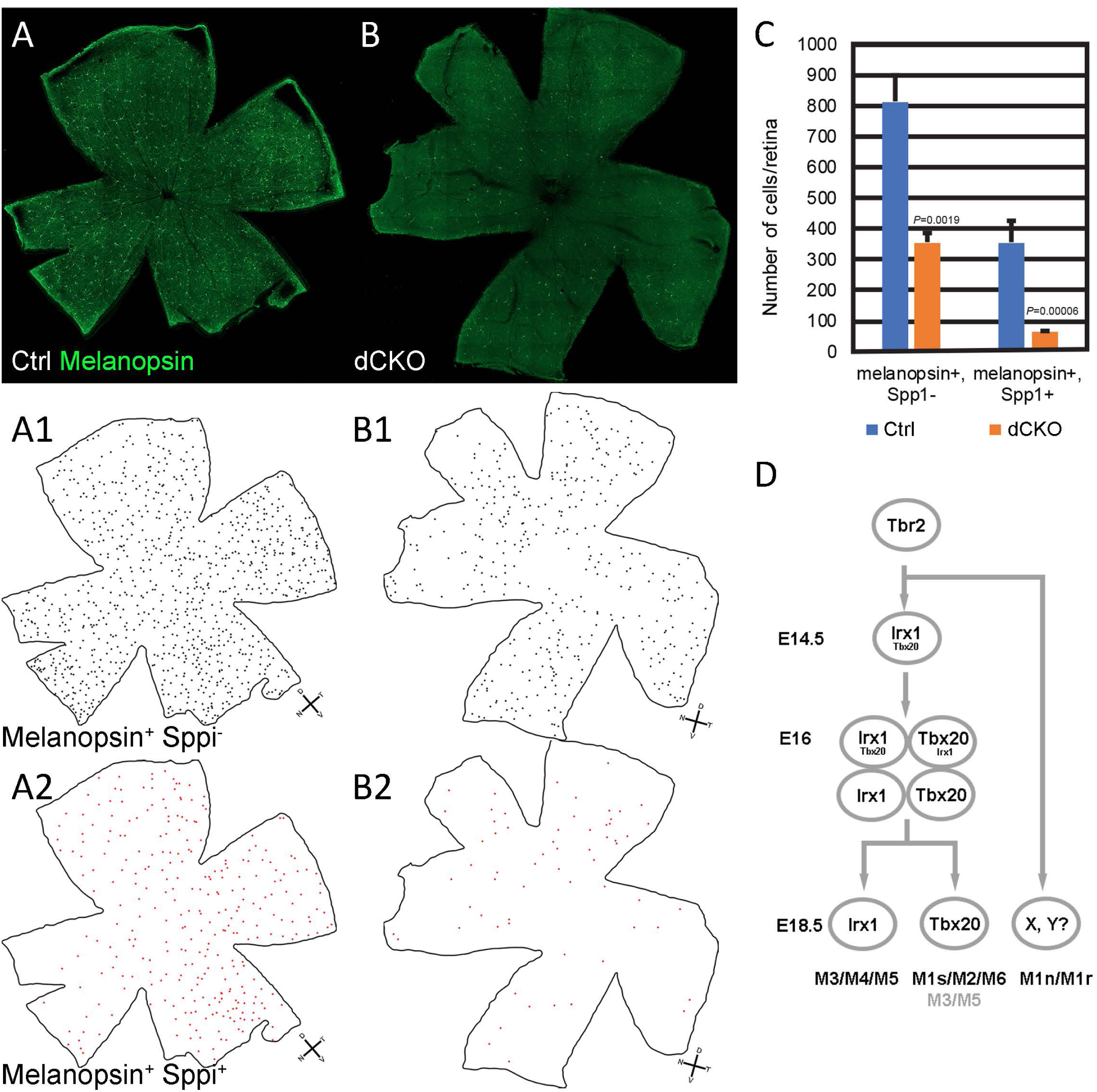
Synergistic role of *Irx1* and *Tbx20* in regulating the formation of ipRGC. **(A, B)** IF staining on *Irx1^f/+^; Tbx20^f/+^* (*Ctrl*) (A) and *Six3-Cre; Irx1^f/f^; Tbx20^f/f^* (*dCKO*) (B) retinal flatmounts using anti-melanopsin and anti-Spp1 antibodies. **(A1, B1)** Lucida tracing images of melanopsin+ Spp1-ipRGCs in A and B. **(A2, B2)** Lucida tracing images of melanopsin+Spp1+ ipRGCs (M2 and M4) in A and B. **(C)** Quantitative RT-PCR analysis of *Opn4* expression between P30 *Ctrl* and *dCKO* retinas. **(D)** A working model illustrating the Tbr2-dependent mechanism of ipRGC subtype lineage segregation.

## DISCUSSION

The emergence of diverse neuronal types in the central nervous system, each with unique morphophysiological characteristics, is a complex process governed by discrete transcription regulatory networks that unfold dynamically across developmental time and space. In the mouse retina, ipRGCs are the first RGCs to emerge, with the onset of Tbr2 expression marking the initiation of this process, ultimately leading to the development of six unique adult ipRGC subtypes. In this study, we identified two major transcription factors downstream of Tbr2, namely Irx1 and Tbx20, which play pivotal roles in regulating *Opn4* expression and the formation of distinct subsets of ipRGCs. These two factors co-exist briefly in E16.5 RGCs, after which they segregate to specify different paths through which distinct ipRGC subtypes emerge.

In mature retina, Irx1 is expressed mainly in M3, M4, and M5 ipRGCs. Loss-of-function study indicates that Irx1 controls *Opn4* expression but is dispensable for the development and maintenance of the ipRGCs it specified. This interesting observation might be rooted in the fact that there are six members of the *Irx* family genes in mice (*Irx1* to *Irx6*) and most are detected in the retina in some RGCs, including ipRGCs, in public scRNA-seq datasets. It is therefore likely that other *Irx* family genes may compensate for *Irx1* in controlling the formation of Irx1-expressing ipRGCs. In the developing mouse cortex, *Tbr2* is a key regulator of intermediate (basal) neuronal progenitor cells^42, 43^. A ChIP-seq study has identified a Tbr2-bound, 570 bp fragment 5868 bp downstream of *Irx1* containing multiple Tbr2/Eomes binding sites (Chr13: 72089243-72089812, mm9) (Extended Figure 14A)^44^, implicating that Tbr2 may also bind to *Irx1* directly in RGCs during early development and that *Tbr2-Irx1* transcription cascade plays a crucial role in orchestrating the proper gene expression program in the developing M3, M4, and M5 ipRGCs.

The expression of Tbx20 in ipRGCs has been reported previously^45^; however, the specific ipRGC subtypes in which it is expressed and its functional role remain unclear. Here we show that Tbx20 is primarily expressed in M1s, M2, and M6 subtypes, with limited expression in subsets of M3 and M5 ipRGCs in adult retina. Tbx20 regulates the formation of these cells and maintains the gene expression programs essential for their survival. These findings resonate well with the general notion that T-box TFs exhibit distinct spatiotemporal expression patterns during diverse developmental processes and typically contribute to both the formation of specific cell types and the regulation of gene expression programs critical for their functions ^46–52^.

Our cell type classification data, based on recordings from distinct TF-driven reporters in this and previous work involving Tbr2, call for an expansion of RGC diversity currently based on scRNA-seq data but with partial reference to morphologically and functionally categorized types. Using the widely studied M1 ipRGC as an example, three sub-groups were described among Tbr2-expressing RGCs—M1s (26%), M1r (28%), and M1n (46%)—based on their distinctive IMP profiles, intrinsic photosensitivity, and spiking patterns^22^. Lee *et al.* independently reported two subtypes of M1 ipRGCs, M1a and M1b, distinguishable by their synaptic connectivity to glutamatergic amacrine cells expressing the vesicular glutamate transporter 3^53^. Although these published M1 sub-groups are noted to be morphologically similar and their dendrites stratify to the junction of INL and IPL, they can nonetheless be discerned by stereotypical IMPs, synaptic connections, and other circuit-level properties. Our study identifies the least abundant IMP subtype M1s to be the major M1 subtype labeled among Tbx20-expressing ipRGCs. This suggests that the specification and development of the more abundant M1r and M1n subtypes rely on transcription factor(s) other than Tbx20 and Irx1.

The M3 and M5 ipRGCs were likewise noted to be diverse, and several unique IMP profiles were described among Tbr2-expressing cells^22^. In the current study, we renamed the three discernible M3 subtypes M3s, M3n, and M3r, corresponding to previously characterized Tbr2-expressing cells (Figure 6h, 6i, and 6j cells in Chen *et al., 2021)*^22^ based on characteristics in their IMP profiles. What’s interesting is that Irx1 did not specify M3s, and Tbx20 did, while both specified M3n and M3r.

Even more interesting is that the novel M5nt (see below) labeled in the *Irx1-Ai9* retinas had readily detectable intrinsic photosensitivity (Figure 3C), while those in *Tbx20-Ai9* retinas were barely detectable (Figure 5H). The phototransduction pathways downstream of melanopsin are mediated by members of the Gq family, but the downstream effectors appear diverse^54–58^, which might account for the sensitivity differences we observed here. We renamed the diverse *Tbr2*-expressing M5 ipRGCs described in Figures 8d, 8e, and 8f of Chen *et al.*^22, 55^ to be M5r, M5s, and M5n, respectively, based on the nomenclature guidelines described earlier. Moreover, we identified a new M5 subtype in both Tbx20- and Irx1-expressing ipRGCs in this study and coined it M5nt (no tail), which lacks the characteristic tails (arrows in Figure 3D and 5G) following the cessation of positive current steps. Interestingly, the chance of finding an M5 ipRGC is greater in Irx1-expressing cells (27%) than in Tbx20-expressing cells (8%), and the majority of the M5 subtype encountered in Irx1-expressing cells is M5nt (28/44, 64%). The M5s subtype previously seen in Tbr2-expressing cells was not labeled in either line, suggesting that transcription factor(s) other than Irx1 and Tbx20 specify its fate.

Taken together, we have revealed a key mechanism by which the Tbr2-mediated gene regulatory network controls ipRGC subtype specification during retinal development. Based on in-depth analysis of scRNA-seq data, *Irx1* is activated first in Tbr2-expressing RGCs around E14 (Extended Figure 14B). At this stage, *Tbx20* is barely detectable, and the two Tbr2+ Irx1+ clusters, C4 and C7, are distantly positioned on the UMAP plot and express little *Opn4*, indicating that these cells have not yet acquired the transcriptomic signature characteristic of ipRGCs (Extended Figure 14C). Subsequently, between E14.5 and E15.5, *Tbx20* expression begins to emerge in a subset of Tbr2+ Irx1+ RGCs. At this point, cells expressing Irx1 and/or Tbx20 have not yet fully committed to an ipRGC fate (Figure 7). By E16.5, while most Tbx20-expressing cells are destined to give rise to adult Tbx20+ ipRGCs, a small fraction of Irx1-expressing cells have not yet locked in their terminal fate. By E18.5, the terminal fates of both Irx1- and Tbx20-expressing cells appear to be fully established. The sequential activation of these genes during development thus marks the initial lineage segregation events that specify ipRGC subtypes. However, their expression does not encompass all known ipRGC subtypes. It is evident that additional factors, likely activated in Tbr2-expressing cells at a later stage, are involved in the formation of other M1 subtypes, such as M1n and M1r (Figure 10D). Reanalysis of scRNA-seq datasets from various stages has indeed identified several candidate transcription regulators absent at E16 but emerging at P0, with their expression restricted to specific M1 and/or M2 ipRGCs in the adult retinas (data not shown).

*Tbr2-Irx1* and *Tbr2-Tbx20* transcriptional cascades operate together transiently in fate-undetermined Tbr2-expressing RGCs between E15.5 and E16.5, and hence it is possible that they interact to specify the formation of a subset of ipRGCs. This notion is supported by the IF data showing that the number of ipRGCs drops more significantly in double knockout (Figure 10A-10C). By E18.5, these two pathways begin to operate independently in distinct ipRGC subtypes. Notably, they both contribute to regulating *Opn4* expression in the ipRGC subtypes they specify, and it will be interesting to see how they might do that. Current studies have only identified Tbr2 as the key regulator not just for the formation of ipRGCs but also crucial for maintaining the expression of *Opn4*^19, 22, 31^. Several upstream regulators of *Tbr2* have been implicated in controlling *Opn4* expression and ipRGC identity, potentially through Tbr2-dependent pathways^27, 32, 59–63^. In addition to TFs that positively regulate the genesis of ipRGCs, some negative regulators have also been implicated. For example, Pou4f1 is a TF known to be expressed in RGCs with a mutually exclusive pattern with ipRGCs^19, 22^. In the absence of *Pou4f1*, *Irx1* has been shown to be upregulated^64^. Our previous study also identified Pou4f1-bound elements near *Irx1* and *Tbx20* loci, implicating that Pou4f1 plays a negative role in ipRGC genesis^30^. It would be of interest to understand whether and how these TFs interact with one another in controlling the development, gene expression, axonal projection, dendritic morphology and synaptic connections, and subtype-specific physiology of distinct ipRGC subtypes.

In summary, we have identified two major downstream effectors of Tbr2 that function primarily independently in different ipRGC subtypes. Both factors play critical roles in regulating key marker genes and contribute to ipRGC subtype specification, either independently or in combination. However, the mechanisms by which naïve Tbr2-expressing RGCs activate these genes to establish subtype identity remain unclear. This transition from a naïve state to a committed cell fate likely involves interactions between transcription factors and epigenomic regulatory elements within developing Tbr2⁺ cells at key developmental time points. Future studies investigating how these transcription factors engage with the genome and how the genome responds during fate determination and subtype specification will be essential for understanding the terminal differentiation and molecular diversity of RGCs, as well as the specific roles these factors play in the process.

## ACKNOWLEDGMENTS

This work was supported by grants from the National Institutes of Health to C.-A.M. (EY024376), C.-K.C (EY032898, EY034219), and B.A.A. (DE028527, DE026433, T90DE023520). This work was also supported in part by National Eye Institute Vision Core Grants P30EY028102 (UTHealth), the Hermann Eye Fund (UTHealth), and the Stephen Lasher Endowed Professorship (UTHealth). We acknowledge the Genetically Engineered Mouse Facility at The University of Texas MD Anderson Cancer Center for making *Irx1^HA3-P2ACreERT2/+^* and *Tbx20^HA3-P2ACreERT2/+^* mouse lines. *Irx1^flox^* mouse was generated by GemPharmatech USA (San Diego, CA; Strain #: T008697). *Tbx20^flox^* was purchased from Jackson Lab (Bar Harbor, ME; Strain #:024665; RRID:IMSR_JAX:024665). *Six3-Cre* was a gift from Dr. Yas Furuta. *Opn4^Cre^* was a gift from Dr. Samer Hattar. *Rosa^iAP^* was a gift from Dr. Tudor Badea. We thank Andrew Kim and Amber Lewis for technical assistance.

## AUTHOR CONTRIBUTIONS

TK, CKC, and CAM conceptualized and designed experiments. TK, CKC, HYA, DS, SE, BAA, LS, YJC, and CAM executed experiments. CAM and CKC wrote the paper.

## DECLARATION OF INTEREST

The authors declare no competing interests.

## METHODS

### Animals

*Irx1^HA3-P2ACreERT2^* and *Tbx20^HA3-P2ACreERT2^* mice were generated by inserting a synthetic HA3-P2A-CreERT2 DNA fragment in the c-terminal end in-frame with the last translation codon using CRISPR/Cas9 knock-in (KI) technique (described in Extended Figure 2. Note: these alleles are functional alleles). *Irx1^flox^* mice were generated by GemPharmatech (Cat no: T008697). Two loxP sites were inserted in *Irx1*’s intron 1 and downstream to 3’ UTR, respectively, using CRISPR/Cas9 KI technique. The exons 2-4 of the *Irx1^flox^* allele would be removed upon Cre-mediated deletion (Figure 8). *Irx1^LacZ^* mouse was described previously.^65^ PCR primers used for genotyping the *Irx1^HA3-P2ACreERT2^*, *Irx1^flox^*, and *Irx1^LacZ^* alleles were listed in Key Resource Table. The generation and genotyping of *Six3-Cre*, *Rosa^iAP^*, *Ai9*, and *Opn4^Cre^* mice were described previously.^25, 66–68^ All mice were maintained on C57BL6/129 mixed backgrounds.

Mouse lines (Irx1*^HA3-P2ACreERT2^*; Rosai*^AP/+^*, Irx1*^HA3-P2ACreERT2^*; Ai9, Tbx20*^HA3-P2ACreERT2^*; *Rosai^AP/+^*, and *Tbx20^HA3-P2ACreERT2^*; *Ai9*) of either sex at various ages between P20 to 6 months were used for sparse labeling or recording/dye filling experiments. Data in Figures 8-10 and Extended Figures 9-13 were obtained using littermate control and mutant mice. Pre-weaning animals were housed with their mother; weaned animals were housed in groups of no more than 5 in individually ventilated cages. All animal procedures followed the US Public Health Service Policy on Humane Care and Use of Laboratory Animals and were approved by the Institutional Animal Care and Use Committee at The University of Texas McGovern Medical School at Houston and the Animal Welfare Committee at The University of Texas Health Science Center at San Antonio.

### Immunohistochemical analysis

Retinal sections or flat-mounted retinas were fixed with 4% paraformaldehyde (PFA) (Electron Microscopy Sciences, Hatfield, PA) and then incubated with the primary antibodies listed in the KEY RESOURCE TABLE below. Alexa Fluor-conjugated secondary antibodies were obtained from Life Technologies (1:800 dilution). DAPI (2 µg/ml) was used to label the nuclei. Images were acquired on Zeiss 800 confocal laser scanning microscope (Carl Zeiss, Thornwood, NY, USA) and exported as TIFF files into Adobe Photoshop (Adobe Systems, San Jose, CA, USA). Cell counting was conducted using the cell counter plugin of NIH ImageJ.

### X-gal staining

X-gal staining was conducted as previously described.^69^ Flat-mounted retinas were fixed in 10% formalin for 30 minutes and washed three times with wash buffer (0.1 M sodium phosphate containing 2 mM MgCl_2_, 0.01% deoxycholate, and 0.02% NP-40). The color reaction was performed in wash buffer containing 5 mM potassium ferrocyanide, 5 mM potassium ferricyanide, and 1 mg/ml X-gal at 25 °C overnight, and terminated in the 10% formalin for 10 minutes. Post-fixed sections were washed, dehydrated, and mounted with Cytoseal 60 (Thermo Scientific, Waltham, MA, USA). Images were collected with a Canon EOS 10 digital camera mounted on an Olympus IX70 microscope (Olympus USA, Center Vally, PA, USA).

### Bulk RNA-sequencing analysis

Twelve retinas from E15.5 *wildtype* and *Six3-Cre; Tbr2^f/f^* embryos from multiple litters were pooled, and RNA was extracted using an RNAeasy mini kit (QIAGEN Science, Germantown, MD, USA). Bulk RNA sequencing was performed in Novogene (Sacramento, CA). Sequencing libraries were prepared using standard Illumina protocol. RNA sequencing was performed using Illumina HiSeq 6000. Illumina Casava was used for base calling. The RNA-seq reads were mapped to the mouse genome (mm10) via STAR (v2.5). HTSeq (v0.6.1) was used to count the read numbers mapped to each gene. Differential expression analysis of two groups was performed using the DESeq2 R package (v2_1.6.3). The resulting P-values were adjusted using Benjamini and Hochberg’s approach for controlling the false discovery rate. Genes with an adjusted P-value < 0.05 were considered as differentially expressed. The raw datasets and normalized count data for each group have been deposited in the NCBI Gene Expression Omnibus repository (GSE149388).

### Single-cell RNA-sequencing data analysis

The single-cell RNA-seg datasets used to profile developing mouse retinal ganglion cells were obtained from the Gene Expression Omnibus (GSE185671).^33^ The raw_count_matrix files were imported into R to create Seurat objects. To ensure data quality, several criteria were applied on a per-cell basis to remove low-quality or dying cells: 1) cells with less than 600 genes, 2) cells with less than 500 and more than 20000 RNAs, and 3) cells with more than 10% of mitochondria RNAs. After filtering out low-quality cells, the data were normalized, scaled, and explored using a standard Seurat workflow with default settings.^70^ Principal component analysis, Louvain algorithm-based clustering, and the UMAP were performed. After the initial clustering, those cells with lower levels of Rbpms expression (Rbpms <1) were removed. The remaining cells were re-clustered. By applying these procedures, 14439 E14 RGCs were divided into 16 clusters, 11810 E16 RGCs were divided into 24 clusters, and 16152 P0 RGCs were divided into 28 clusters.

### Tracing RGC axon targets

For inspecting *Irx1^CreERT2^* or *Tbx20^CreERT2^* driven *Rosa^iAP^* expression in brains, 1 µl of 4-hydroxytamoxifen (4-OHT) was injected into the vitreous of the right eye of *Irx1^CreERT2^*; *Rosa^iAP^* or *Tbx20^CreERT2^*; *Rosa^iAP^* mice using a 33-gauge NanoFil system (World Precision Instruments, Sarasota, FL). Two weeks after injection, animals were anesthetized and perfused with 4% PFA. Whole brains were dissected and post-fixed with 4% PFA. Cryopreserved brains were sectioned into 100 µm, consecutive coronal sections for alkaline phosphatase (AP) staining.

### Alkaline phosphatase (AP) staining

Tamoxifen-activated *Irx1^CreERT2^*; *Rosa^iAP^ or Tbx20^CreERT2^*; *Rosa^iAP^* (Figures 2 and 4) mice were used for AP staining, which was conducted as previously described with minor modifications^71, 72^. Whole eyeballs were fixed with 10% formalin for 10 min. The retinas were removed and flat-mounted on a piece of nitrocellulose membrane, post-fixed for 5 min at room temperature in 10% formalin, washed twice in PBS+ (PBS plus 2 mM MgCl_2_), and heated in PBS for 30 min in a 65°C water bath to inactivate endogenous AP activity. AP staining was performed in 0.1 M Tris (pH 9.5), 0.1 M NaCl, 50 mM MgCl_2_, 0.34 g/ml NBT, and 0.175 g/ml BCIP solution overnight at room temperature with gentle shaking. After staining, tissues were washed 3 times for 20 min in PBS, dehydrated through an ethanol series, and then cleared with 2:1 benzyl benzoate/benzyl alcohol. Images were acquired on a Zeiss Axio Imager 2 microscope equipped with a motorized Z drive (Zeiss United States, Thornwood, NY, USA).

### RNAScope *in situ* hybridization (ISH)

ISH was performed using RNAscope technology with minor modifications (Advanced Cell Diagnostics, Newark, CA, USA). Briefly, 9 µm paraffin sections mounted on superfrost-plus glass slides were subjected to RNAscope 2.5 HD Detection Reagents Brown kit or RNAscope Fluorescent Multiplex kit following manufacturer’s protocols. The procedure involved a 5 min simmering in antigen retrieval reagents followed by RNAscope protease III for 30 min at 40°C. After washing 2X in ddH_2_O, the sections were hybridized with RNAscope *in situ* probes for 2 h at 40°C and processed according to the manufacturer’s protocols. The brown signal was examined and collected using an Olympus IX-70 inverted microscope, and the fluorescent signal was visualized and captured using a Zeiss LSM800 confocal microscope. The probes used were mouse Irx1-C1, mouse Tbx20-C1, RNAscope 3-plex positive control probe, and RNAscope 3-plex negative control probe.

### Quantitative Reverse transcriptase PCR (qRT-PCR) analysis

Total RNAs were pooled from 2 retinas isolated from wildtype and mutant littermate mice at different postnatal stages. RNAs were extracted using TRI reagent (Sigma, St. Louis, MO, USA). First-strand cDNA was synthesized using the Superscript III First-Strand Synthesis System. Quantitative PCR was conducted in a CFX Connect Real-Time System (BioRad, Hercules, CA, USA) with iTaq Universal SYBR Green Supermix (#1725122, BioRad).

### Patch Clamp Recording/Filling of Irx1- and Tbx2-expressing RGCs

Whole-cell patch clamp recording was used, as described previously^22, 73^, to obtain intrinsic membrane properties and test the intrinsic photosensitivity of Irx1+ and Tbx20^+^ cells. Retinas from tamoxifen-treated *Irx1^CreERT2/+^; Ai9* or *Tbx20^CreERT2/+^; Ai9* mice were flat-mounted with ganglion cell side up under infrared illumination with the aid of night vision devices in a perfusion chamber on an Olympus BX51WI fixed-stage microscope (Olympus USA) and supplied (3 ml/minute) with a warm (~34°C) carbogenated mammalian Ringer solution containing (in mM) 120 NaCl, 5 KCl, 25 NaHCO_3_, 0.8 Na_2_HPO_4_, 0.1 NaH_2_PO_4_, 2 CaCl_2_, 1 MgSO_4_, 10 D-glucose, 0.01 L-AP4, 0.01 AP5 and 0.04 NBQX. Retinal neurons were visualized by differential interference contrast under infrared illumination (> 900 nm) and Irx1+ or Tbx20+ cells marked by tdTomato expression were identified by brief 50 msec light pulses (35000 R*/rod/sec) delivered and imaged through the mCherry filter set (Olympus USA). The cumulative total exposure of the retina was less than 1 second to preserve light sensitivity. Once a cell was selected, a 20-minute wait time was used to allow dark adaptation before recording. Patch pipettes made from borosilicate glass tubes (8-12 MΩ) were filled with a potassium-based internal solution containing (in mM) 125 K-gluconate, 8 NaCl, 4 ATP-Mg, 0.5 Na-GTP, 5 EGTA, 10 HEPES and 0.2% biocytin (w/v) with pH adjusted to 7.3 using KOH. The liquid junction potential was 15 mV and not corrected. Current clamp recording was conducted using the AM2400 amplifier (A-M Systems, Sequim, WA, USA) driven by the WinWCP program provided by John Dempster (University of Strathclyde, Glasgow, UK). Signals were filtered at 5 kHz and sampled at 10 kHz. To obtain intrinsic membrane property (IMP), a small negative current was injected immediately after break-in to adjust membrane potential from rest to −64 ± 2 mV. This was followed by five 600-msec serial negative current injections for hyperpolarization and four serial positive current injections for depolarization. The current that hyperpolarizes a cell to 100 ± 5 mV was empirically determined and used to adjust the amplitudes of subsequent serial current steps. Under these conditions, cells exhibit characteristic IMP profiles that correspond to unique dendritic morphologies. Intrinsic photosensitivity was initiated by a 15-sec 470 nm light stimulation (~430 μm in diameter) at intensities of 2.67 to 6.14 log R*/rod/sec. Dendritic morphology was recovered *post hoc* by staining for biocytin, and virtually reconstructed in the Neurolucida 360 program (MBF Bioscience, Boston, MA, USA).

### Quantitative Analysis and Statistics

Three pairs of P30 wildtype and littermate mutant mice were used for RT-qPCR (Figures 8 and 9) or cell counting (Figures 2,4, and 10). All data are presented as mean ± standard deviation for each genotype. For comparing the number of ipRGCs (Figure 10), a two-tailed one-sample student’s *t*-test was used. Results were considered significant when *P*<0.05. Statistical tests were conducted using Excel (Microsoft, Redmond, WA, USA).

## KEY RESOURCES TABLE

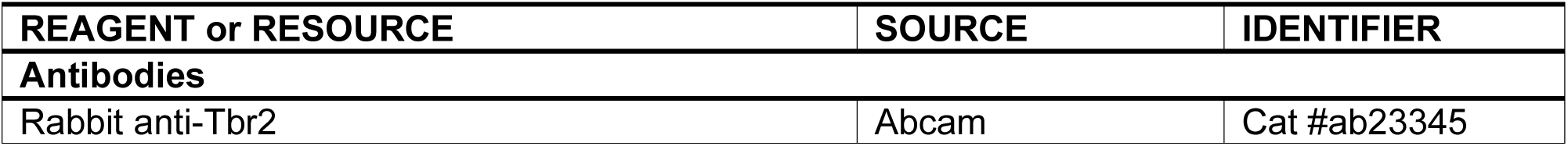
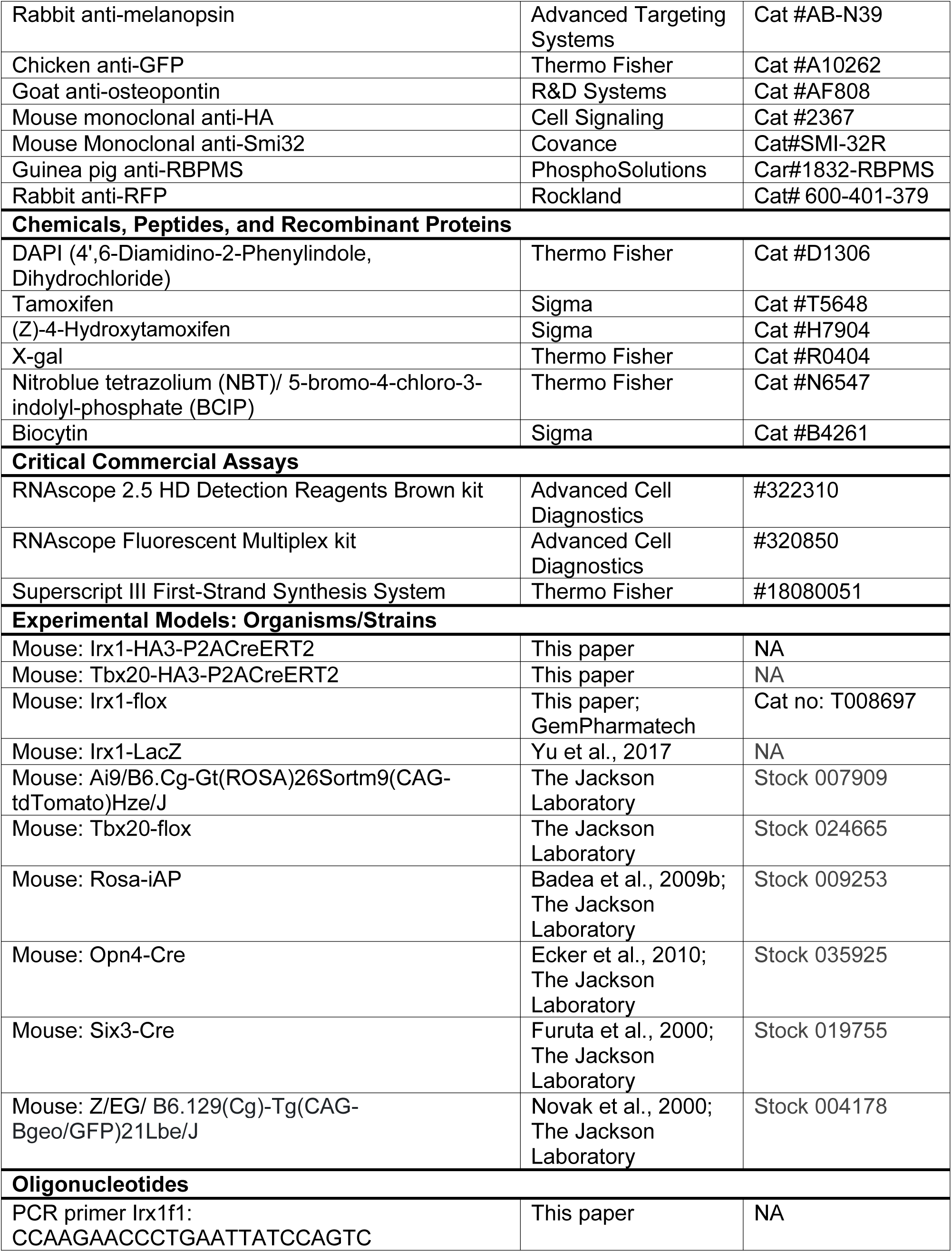
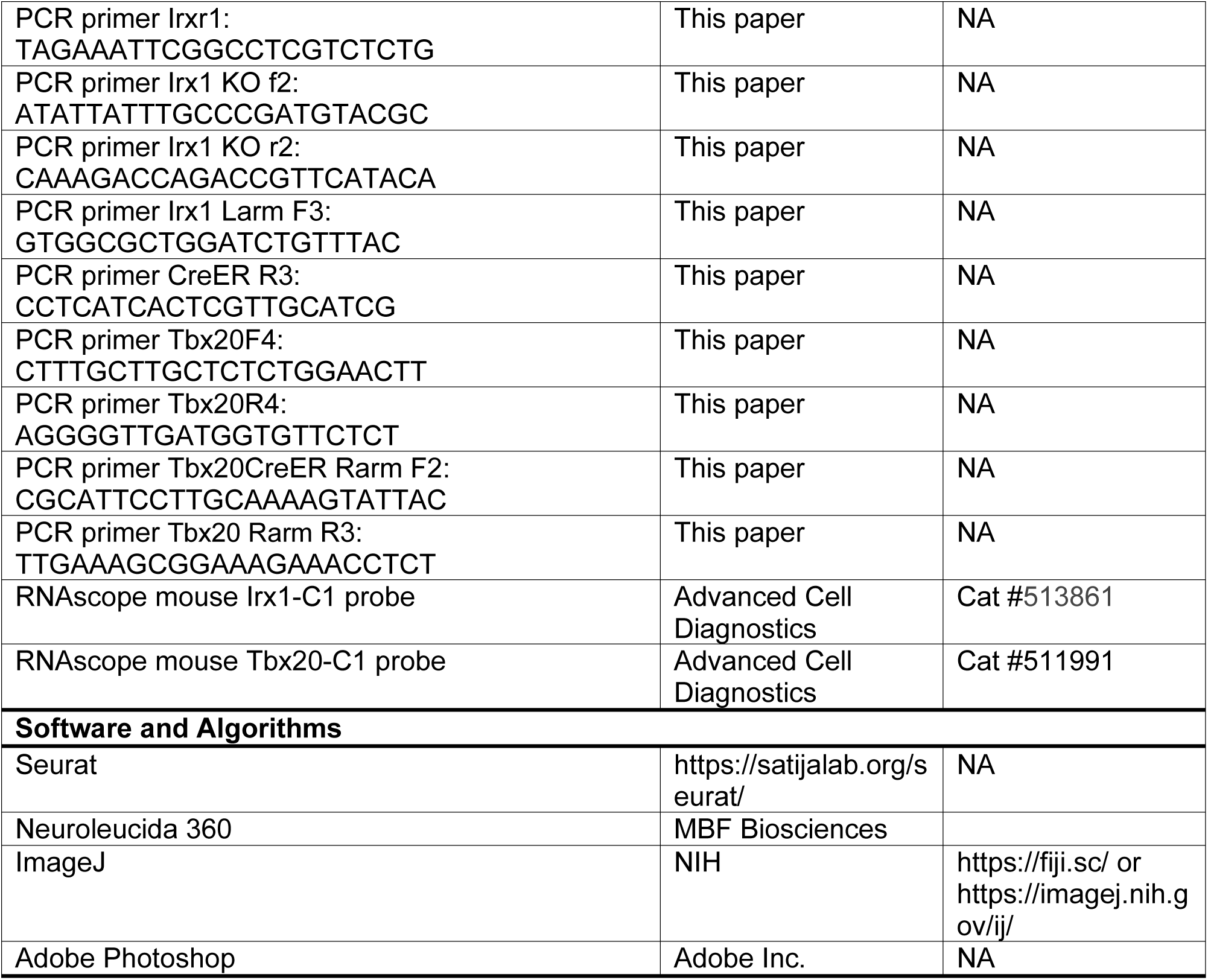

## EXTENDED FIGURE LEGENDS

**Figure 1. Reanalysis of E16 and P0 single cell-RNA sequencing datasets (Geo: GSE185671). (A)** A dot plot showing the 24 clusters identified in the E16 RGC-enriched dataset. **(B)** A dot plot showing the 28 clusters identified in the P0 RGC-enriched dataset. **(C)** A dot plot showing the 5 *Eomes*- and *RBPMS*-enriched clusters using P0 dataset. **(D)** A UMAP plot showing the 28 UMAP clusters of P0 RGCs shown in B. **(E)** Feature heatmap showing the expression of *Eomes*, *Opn4*, *Irx1*, *Tbx20*, *Pou4f1*, and *RBPMS* in P0 RGC clusters.

**Figure 2. Generation of *Irx1^HA3-P2ACreERT2^* and *Tbx20^HA3-P2ACreERT2^* mouse lines. (A)** The genomic structure of *Irx1*, CRISPR/Cas9 targeting strategy, and *Irx1^HA3-P2ACreERT2^* targeted allele. **(B)** The genomic structure of *Tbx20*, CRISPR/Cas9 targeting strategy, and *Tbx20^HA3-P2ACreERT2^* targeted allele.

**Figure 3. Genetically-activated AP staining on *Irx1^CreERT2^; Rosa^iAP^* retinas. (A)** Schematic illustration showing genetically-activated *Rosa^iAP^* reporter by *Irx1^CreERT2^*. **(B-G)** Representative AP stained RGCs on flat-mounted *Irx1^CreERT2^; Rosa^iAP^* retina with low levels of tamoxifen exposure.

**Figure 4. Genetically-activated tdTomato reporter on *Irx1^CreERT2^; Ai9* retinas. (A)** Schematic illustration showing genetically-activated *Rosa-tdTomato* reporter by *Irx1^CreERT2^*. **(B)** Representative tdTomato signals on tamoxifen-activated *Irx1^CreERT2^; Ai9* retina. **(C, D)** IF staining of *Irx1^CreERT2^; Ai9* retina with anti-HA and anti-RFP antibodies (C), and anti-Spp1 antibody (D). Arrowheads indicate the tdTomato+Spp1+ M4 ipRGCs.

**Figure 5. Genetically-activated AP staining on *Tbx20^CreERT2^; Rosa^iAP^* retinas. (A)** Schematic illustration showing genetically-activated *Rosa^iAP^* reporter by *Tbx20^CreERT2^*. **(B, C)** Representative AP staining patterns on flat-mounted *Tbx20^CreERT2^; Rosa^iAP^* retina with low levels of tamoxifen exposure.

**Figure 6. Genetically-activated tdTomato reporter on *Tbx20^CreERT2^; Ai9* retinas. (A)** Schematic illustration showing genetically-activated *Rosa-tdTomato* reporter by *Irx1^CreERT2^*. **(B)** Representative tdTomato signals on tamoxifen-activated *Tbx20^CreERT2^; Ai9* retina. **(C)** IF staining of *Tbx20^CreERT2^; Ai9* retina with anti-HA and anti-RFP antibodies.

**Figure 7. *Tbx20^LacZ^* allele as a proxy for Tbx20 expression (A)** The Genomic structure of *Tbx20 WT*, *Tbx20^flox^*, and *Tbx20^LacZ^* alleles. **(B, C)** IF staining using anti-LacZ and anti-HA antibodies on *Tbx20^HA/LacZ^* retina. **(D)** Lucida tracing of LacZ- and HA-marked Tbx20+ RGCs in *Tbx20^HA/LacZ^* retina showing that more than 98% of HA+ signal can be detected within LacZ+ cells. Scale bar: 50 μm.

**Figure 8. Segregation of Tbx20- and Irx1-expressing RGCs in adult retina. (A, B)** Representative IF staining images using anti-LacZ and anti-HA antibodies on *Tbx20^LacZ/+^*; *Irx1^HA/+^* retina. Scale bar: 50 μm.

**Figure 9. Irx1 controls the expression of *Opn4* in Irx1-expressing ipRGCs.** IF staining of P30 *Irx1^fx/+^* and *Six3-Cre; Irx1^fx/fx^* retinas using an anti-melanopsin antibody showing a drastic reduction of melanopsin intensity in the ON and OFF sublaminae across all retina quadrants. The ON-layer images are orthogonal Z-projections of confocal images focused on the ON sublaminae, and the OFF-layer images are orthogonal Z-projections of confocal images focused on the OFF sublaminae. Scale bar: 100 μm.

**Figure 10. Irx1 controls the expression of *Opn4* in Irx1-expressing ipRGCs.** IF staining of P14 *Irx1^LacZ/+^* and *Opn4^Cre/+^; Irx1^LacZ/fx^* retinas using an anti-melanopsin antibody showing a drastic reduction of melanopsin intensity in both the ON and OFF sublaminae across all retina quadrants. The images are orthogonal Z-projections of confocal images. Scale bar: 100 μm.

**Figure 11. Irx1 is not essential for the formation of Irx1-expressing ipRGCs. (A, B)** X-gal staining of P30 *Irx1^LacZ/+^* (A) and *Six3-Cre; Irx1^LacZ/fx^* (B) retinal flatmounts. LacZ+ cells in the central retinas were used for comparison because *Six3-Cre* works more efficiently in the central region.

**Figure 12. Tbx20 is essential for the formation of a subset of Tbx20-expressing ipRGCs. (A, B)** IF staining using anti-LacZ antibody on P30 *Six3-Cre; Tbx20^f/+^* (A) and *Six3-Cre; Tbx20^f/f^* (B) retinal flatmounts. **(C)** Numbers of LacZ+ cells in A and B.

**Figure 13. Synergistic role of Irx1 and Tbx20 in the formation of ipRGCs.** IF staining on P30 *Irx1^f/+^*; *Tbx20^f/+^* and *Six3-Cre; Irx1^f/f^*; *Tbx20^f/f^* retinas using anti-melanopsin antibody showing a drastic reduction of the number of melanopsin+ cells and the intensity of IF signals in the ON and OFF sublaminae across all retina quadrants. The ON-layer images are orthogonal Z-projections of confocal images focused on the ON sublaminae, and the OFF-layer images are orthogonal Z-projections of confocal images focused on the OFF sublaminae. Scale bar: 100 μm.

**Figure 14. *Tbr2-Irx1* transcriptional cascade in E14 RGCs. (A)** Tbr2-bound fragment near *Irx1* in E14 cortical neurons. The putative Tbr2-bound sites are predicted by JASPAR (jaspar2020.genereg.net/analysis). **(B)** ChIP-qPCR showing Tbr2 binds to the putative enhancer in A. **(C)** A dot plot showing the co-expression of *Tbr2* and *Irx1* in E14 RGCs. **(D)** A dim plot showing the distant relationship between the two Tbr2+ Irx1+ clusters, C4 and C7, in C.

## Notes

**Disclosures:** The authors declare no conflicts of interest

### Competing Interest Statement

The authors have declared no competing interest.

