## Supplemental Figures for "Tbr2-Dependent Parallel Pathways Regulate the Development of Distinct ipRGC Subtypes"

Fig. S1

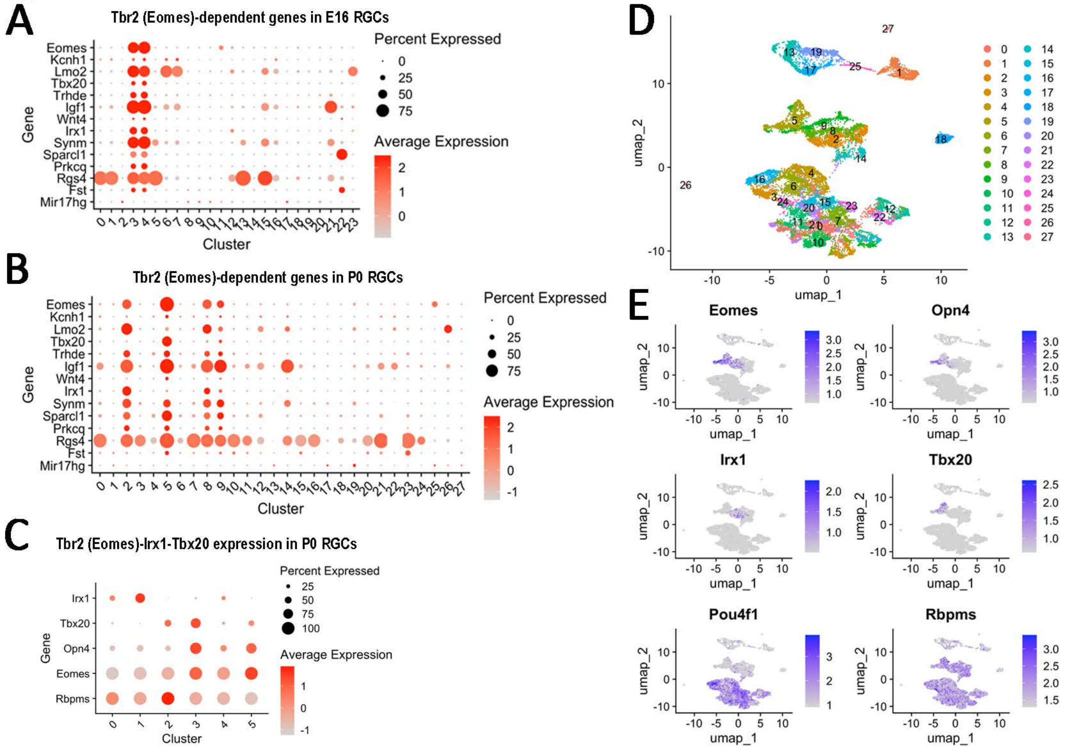
A
Tbr2 (Eomes)-dependent genes in E16 RGCs
Percent Expressed
Average Expression
Gene
Cluster
Eomes
Kcnh1
Lmo2
Tbx20
Trhde
Igf1
Wnt4
Irx1
Synm
Sparcl1
Prkcq
Rgs4
Fst
Mir17hg
B
Tbr2 (Eomes)-dependent genes in P0 RGCs
C
Tbr2 (Eomes)-Irx1-Tbx20 expression in P0 RGCs
Opn4
Rbpms
D
umap_1
umap_2
E
Eomes
Opn4
Irx1
Tbx20
Pou4f1
Rbpms

Fig. S2

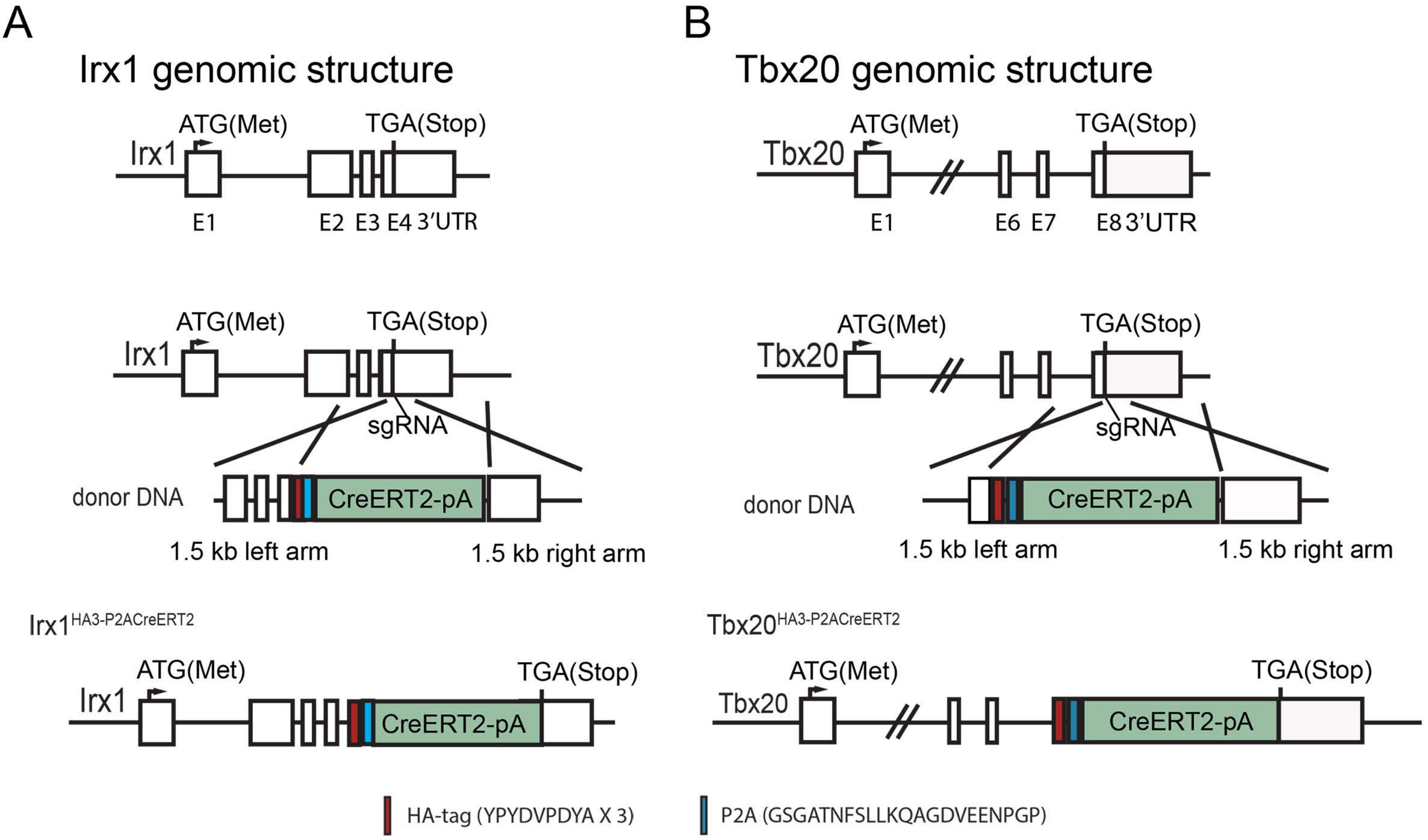

A
Irx1 genomic structure
ATG(Met)
TGA(Stop)
Irx1
E1
E2 E3 E4 3'UTR
ATG(Met)
TGA(Stop)
Irx1
sgRNA
donor DNA
CreERT2-pA
1.5 kb left arm
1.5 kb right arm
Irx1HA3-P2ACreERT2
ATG(Met)
TGA(Stop)
Irx1
CreERT2-pA
B
Tbx20 genomic structure
ATG(Met)
TGA(Stop)
Tbx20
E1
E6 E7
E8 3'UTR
ATG(Met)
TGA(Stop)
Tbx20
sgRNA
donor DNA
CreERT2-pA
1.5 kb left arm
1.5 kb right arm
Tbx20HA3-P2ACreERT2
ATG(Met)
TGA(Stop)
Tbx20
CreERT2-pA
HA-tag (YPYDVPDYA X 3)
P2A (GSGATNFSLLKQAGDVEENPGP)

Fig. S3

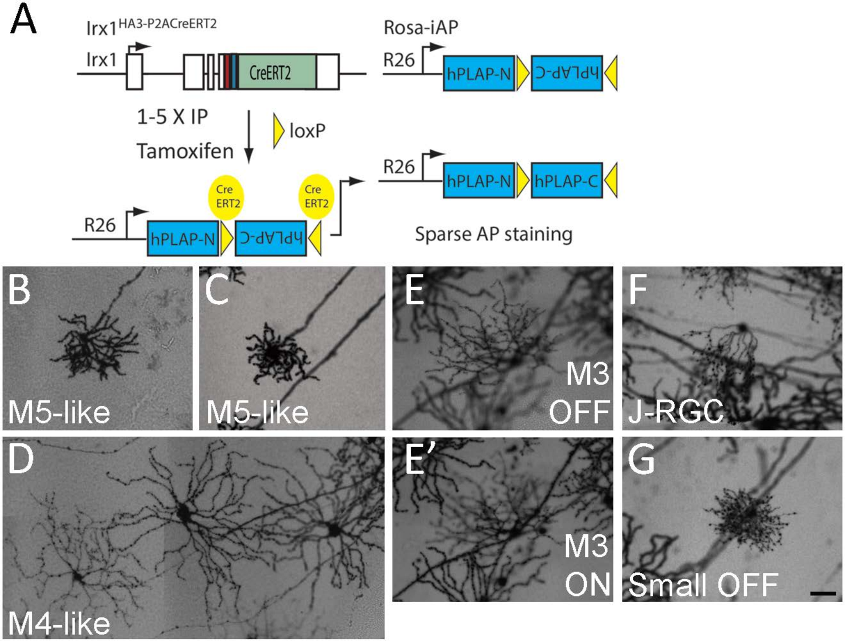
A
Irx1HA3-P2ACreERT2
Irx1
CreERT2
Rosa-iAP
R26
hPLAP-N
hPLAP-C
1-5 X IP
Tamoxifen
loxP
Cre ERT2
Cre ERT2
R26
hPLAP-N
hPLAP-C
R26
hPLAP-N
hPLAP-C
Sparse AP staining
B
M5-like
C
M5-like
E
M3 OFF
F
J-RGC
D
M4-like
E'
M3 ON
G
Small OFF

Fig. S4

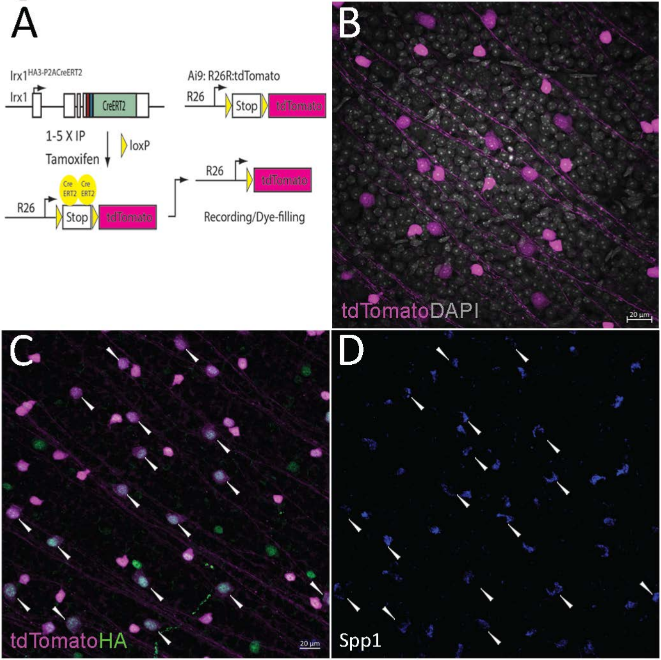
A
Irx1HA3-P2ACreERT2
Irx1
CreERT2
1-5 X IP
Tamoxifen
loxP
Ai9: R26R:tdTomato
R26
Stop
tdTomato
Cre ERT2
Recording/Dye-filling
B
tdTomatoDAPI
20 µm
C
tdTomatoHA
D
Spp1

Fig. S5

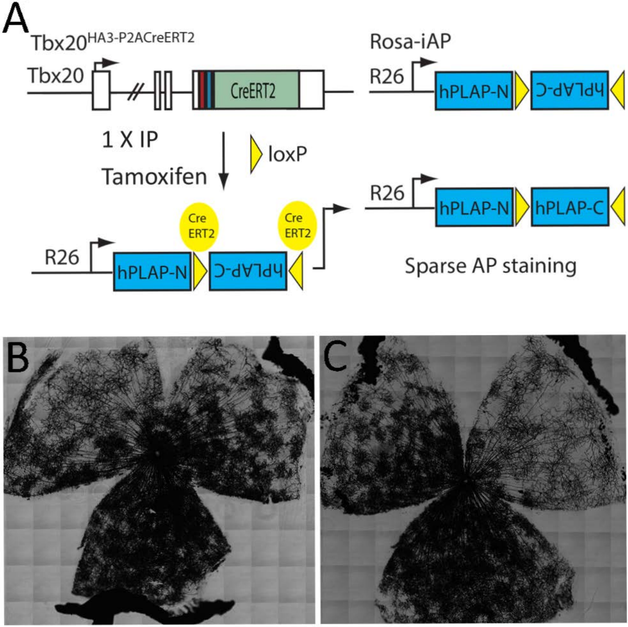
A
Tbx20HA3-P2ACreERT2
Tbx20
CreERT2
Rosa-iAP
R26
hPLAP-N
hPLAP-C
1 X IP
Tamoxifen
loxP
Cre ERT2
Cre ERT2
R26
hPLAP-N
hPLAP-C
R26
hPLAP-N
hPLAP-C
Sparse AP staining
B
C

Fig. S6

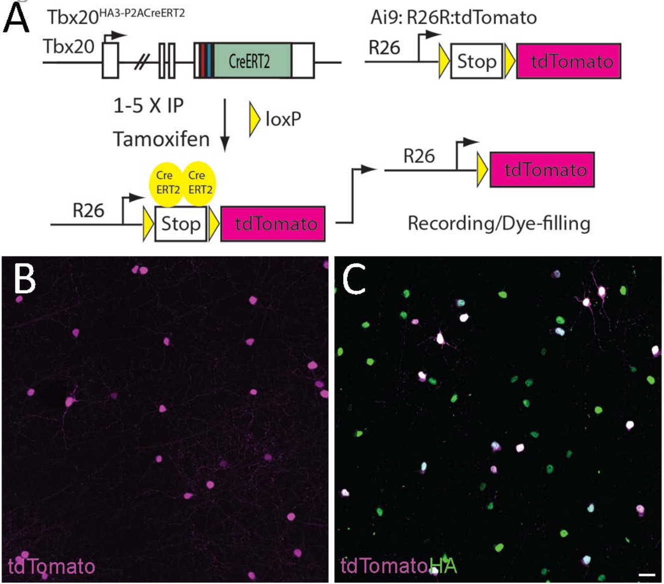

A
Tbx20HA3-P2ACreERT2
Tbx20
CreERT2
Ai9: R26R:tdTomato
R26
Stop
tdTomato
1-5 X IP
Tamoxifen
loxP
Cre ERT2
Cre ERT2
R26
Stop
tdTomato
R26
tdTomato
Recording/Dye-filling
B
tdTomato
C
tdTomato HA

Fig. S7

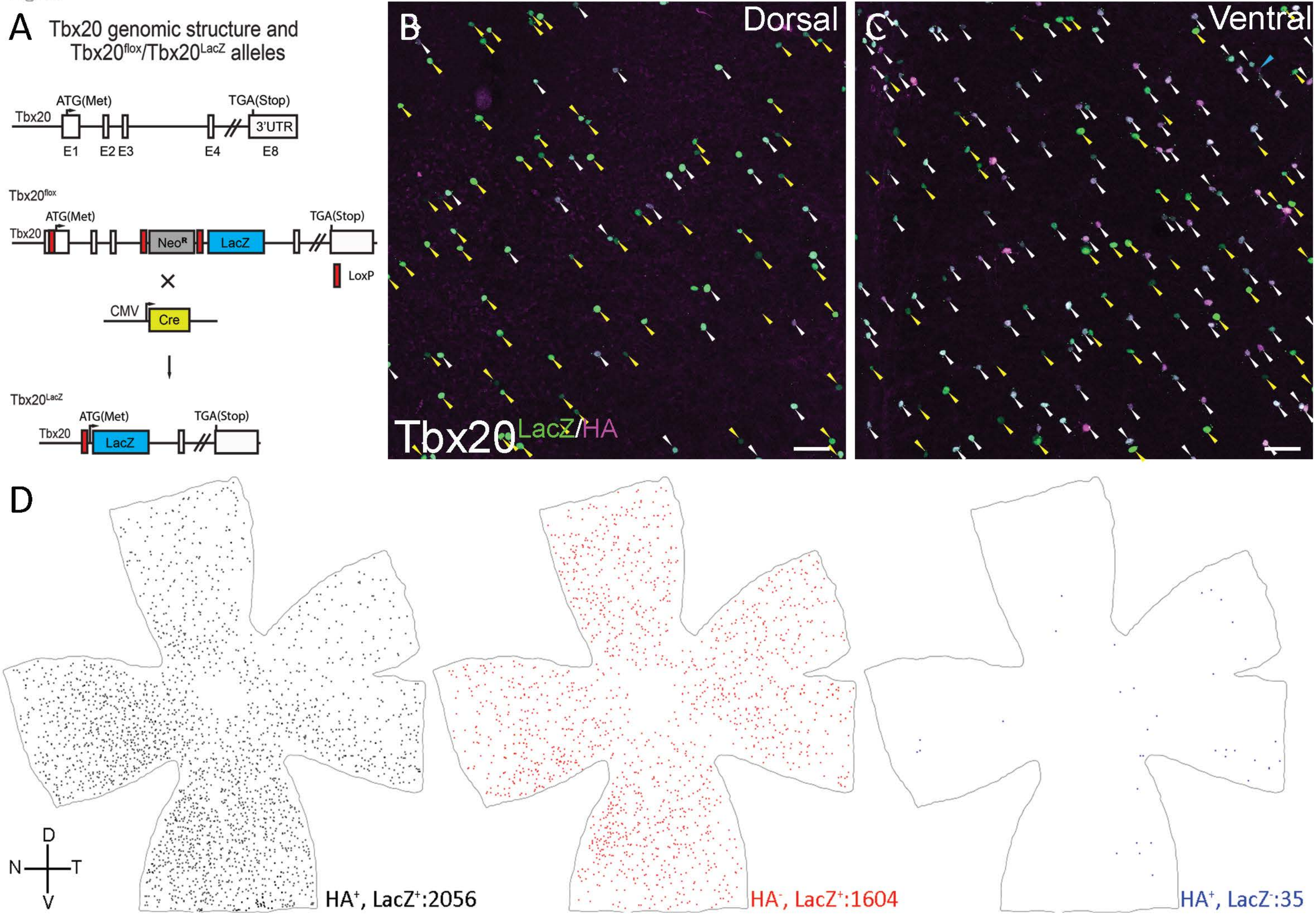
A
Tbx20 genomic structure and
Tbx20flox/Tbx20LacZ alleles
Tbx20
ATG(Met)
TGA(Stop)
3'UTR
E1
E2 E3
E4
E8
Tbx20flox
NeoR
LacZ
LoxP
CMV
Cre
Tbx20LacZ
B
Dorsal
C
Ventral
Tbx20LacZ/HA
D
HA+, LacZ+:2056
HA-, LacZ+:1604
HA+, LacZ-:35
D
N
T
V

Fig. S8

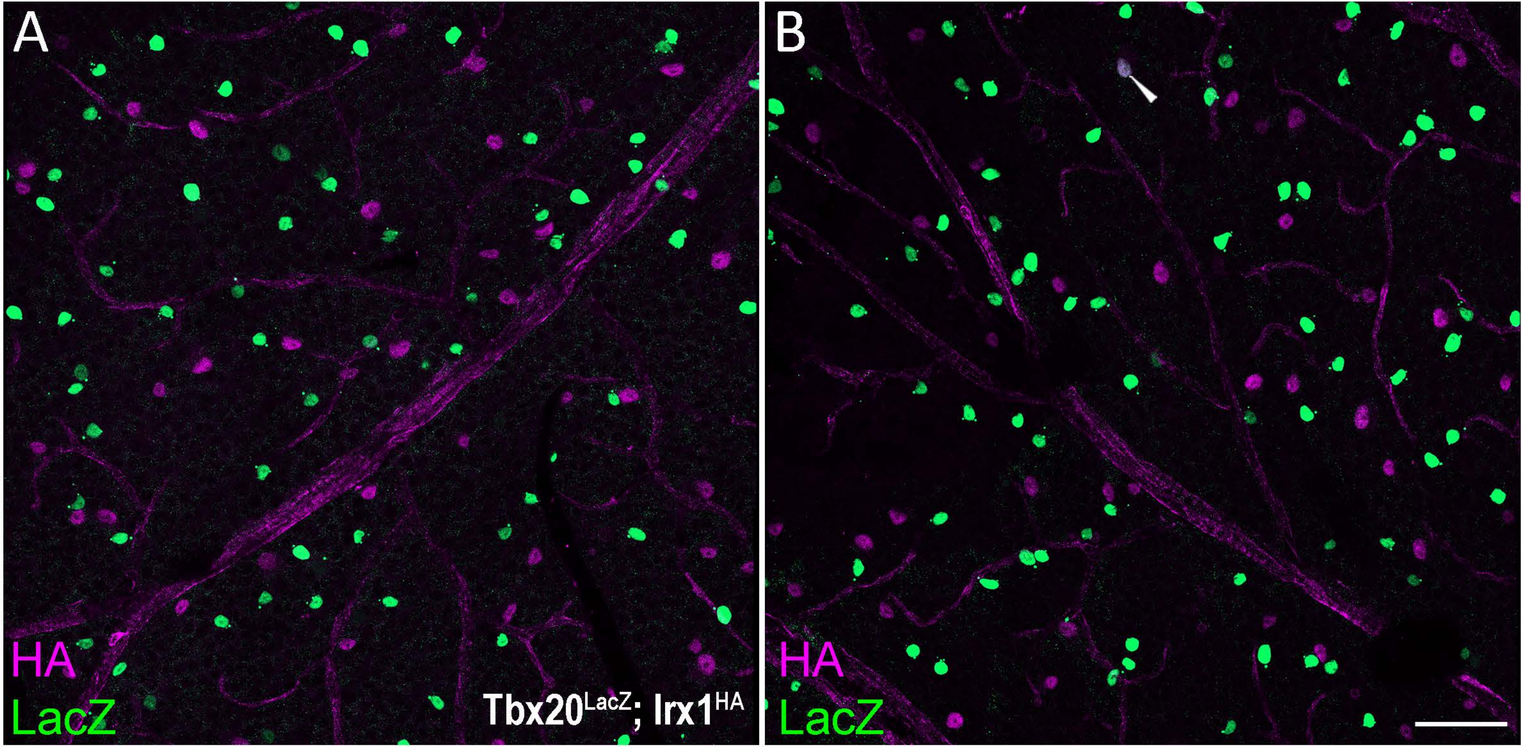

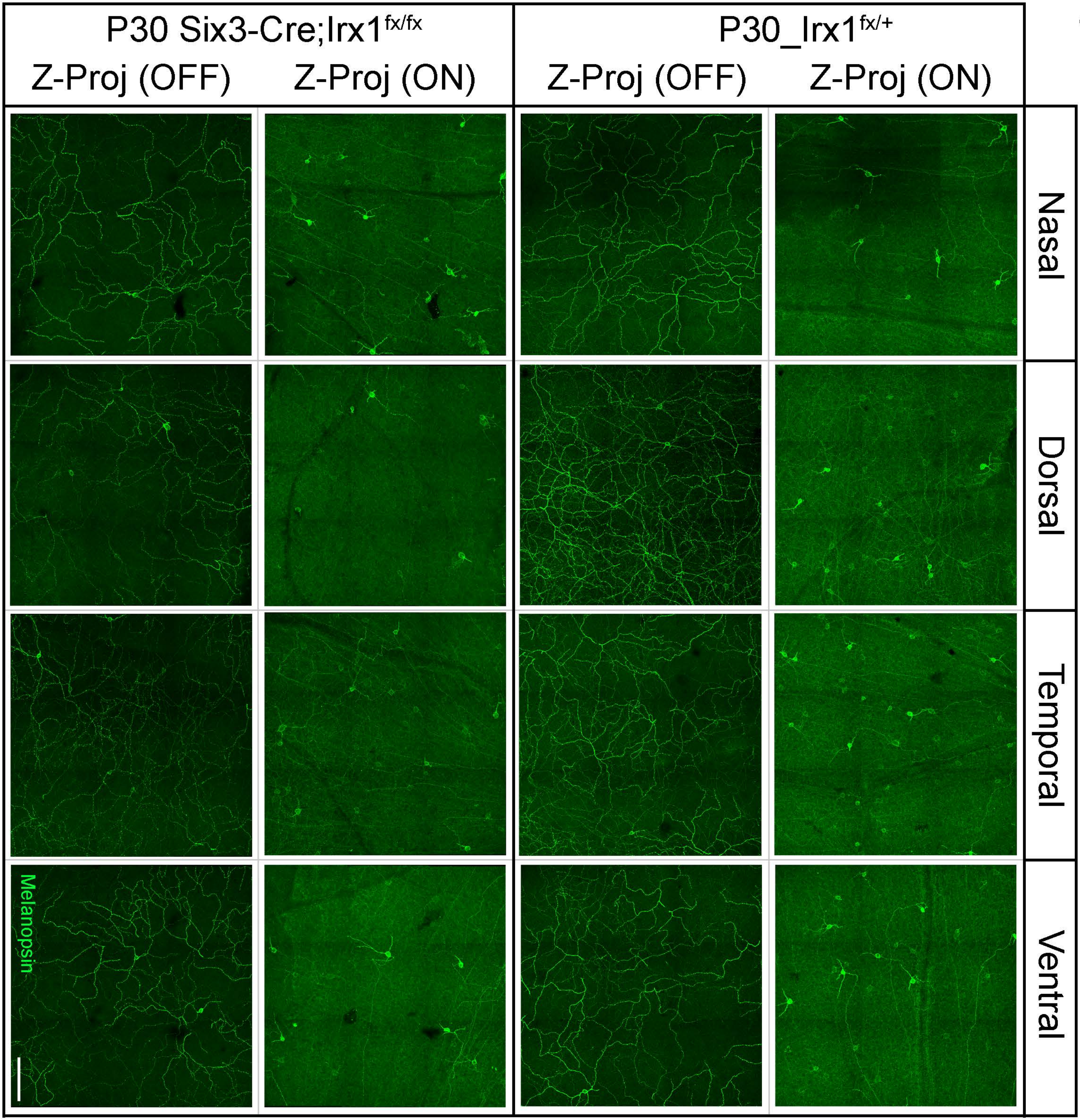
P30 Six3-Cre;Irx1fx/fx
P30_Irx1fx/+
Z-Proj (OFF)
Z-Proj (ON)
Z-Proj (OFF)
Z-Proj (ON)
Nasal
Dorsal
Temporal
Ventral
Melanopsin
Fig. S9

Fig. S10

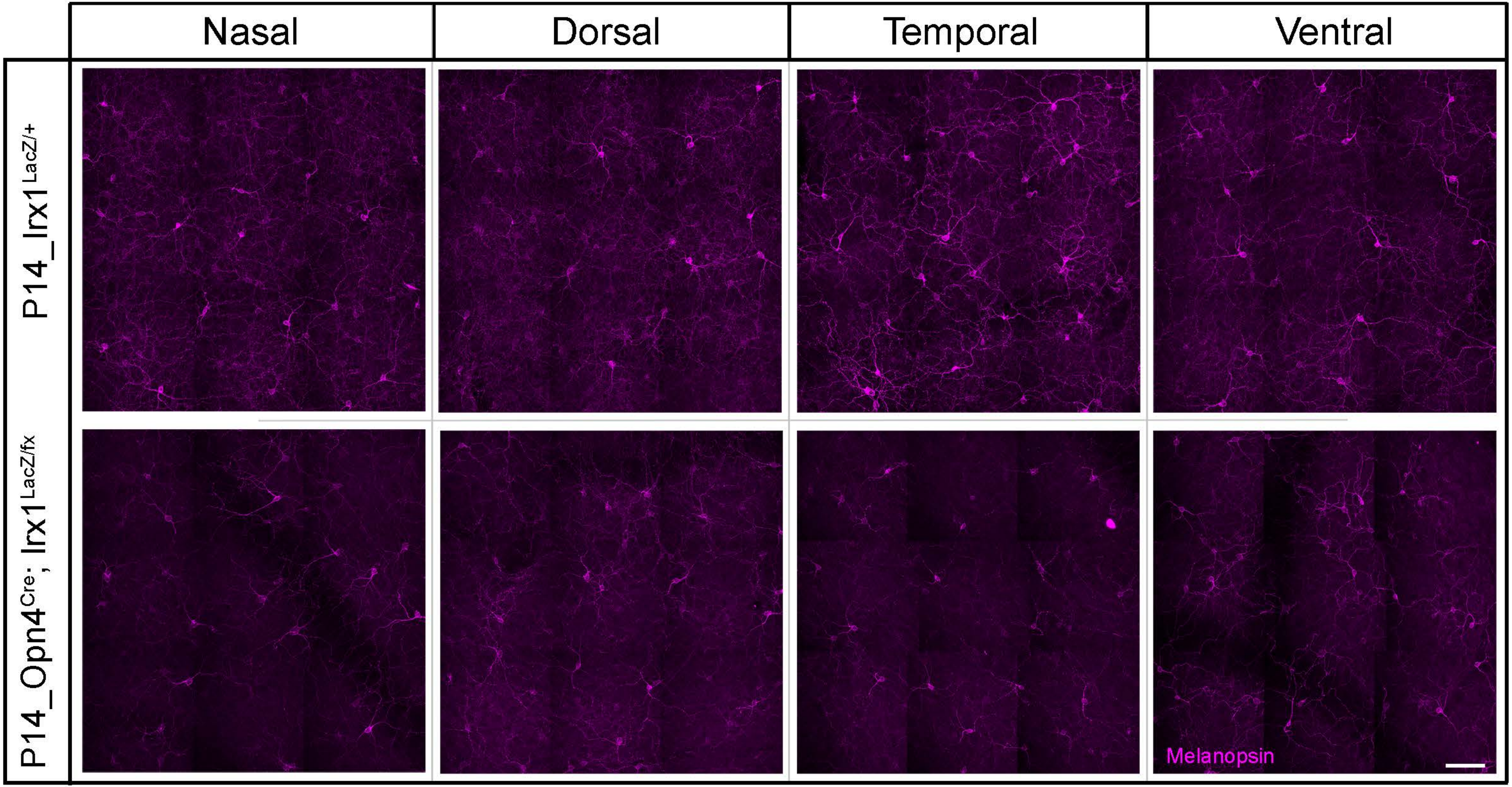

Nasal
Dorsal
Temporal
Ventral
P14_Irx1 LacZ/+
P14_Opn4 Cre/+; Irx1 LacZ/fx
Melanopsin

Fig. S11

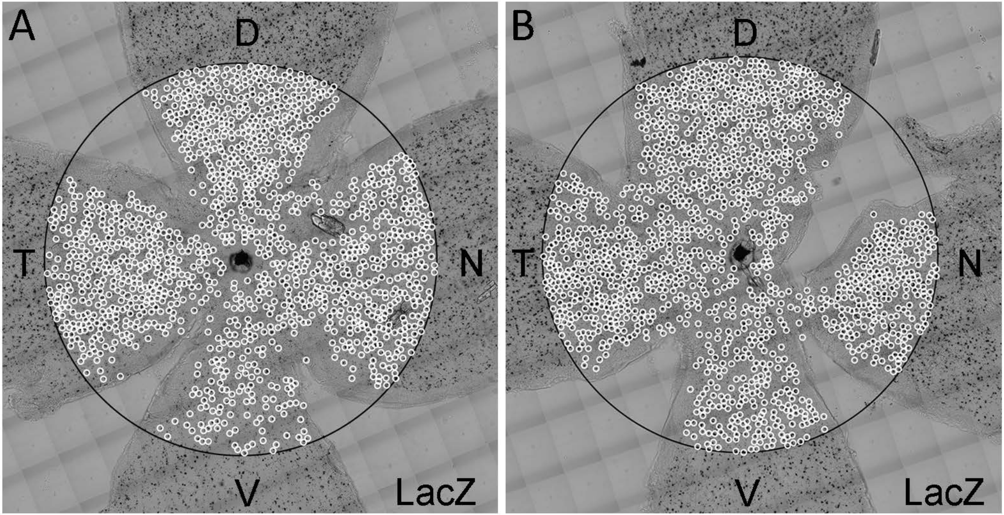


Irx1$^{LacZ/+}$
Central: 1627

Six3-Cre; Irx1$^{LacZ/fx}$
Central: 1690

Fig. S12

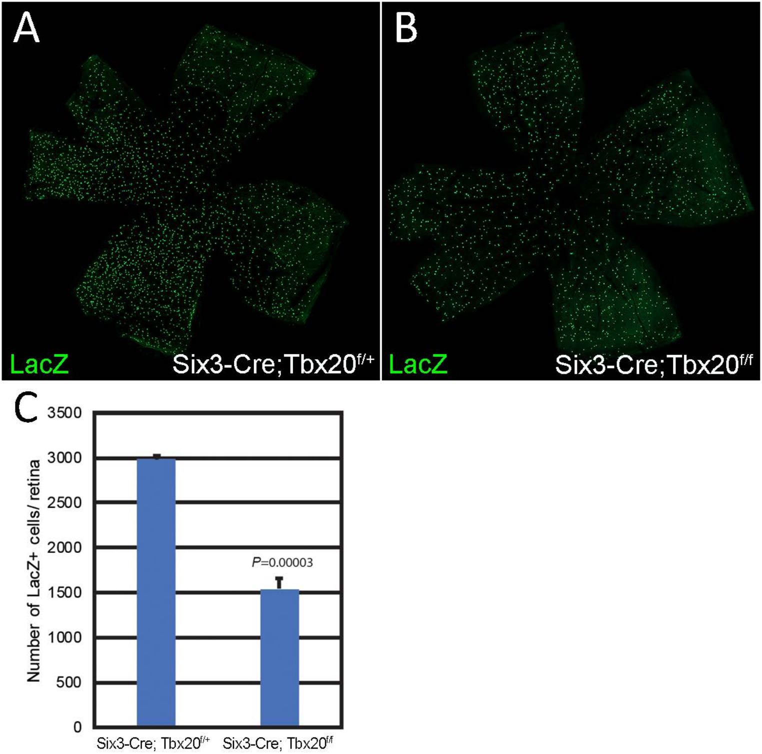

A
B
LacZ
Six3-Cre;Tbx20f/+
LacZ
Six3-Cre;Tbx20f/f
C
3500
3000
2500
2000
1500
1000
500
0
Number of LacZ+ cells/ retina
P=0.00003
Six3-Cre; Tbx20f/+
Six3-Cre; Tbx20f/f

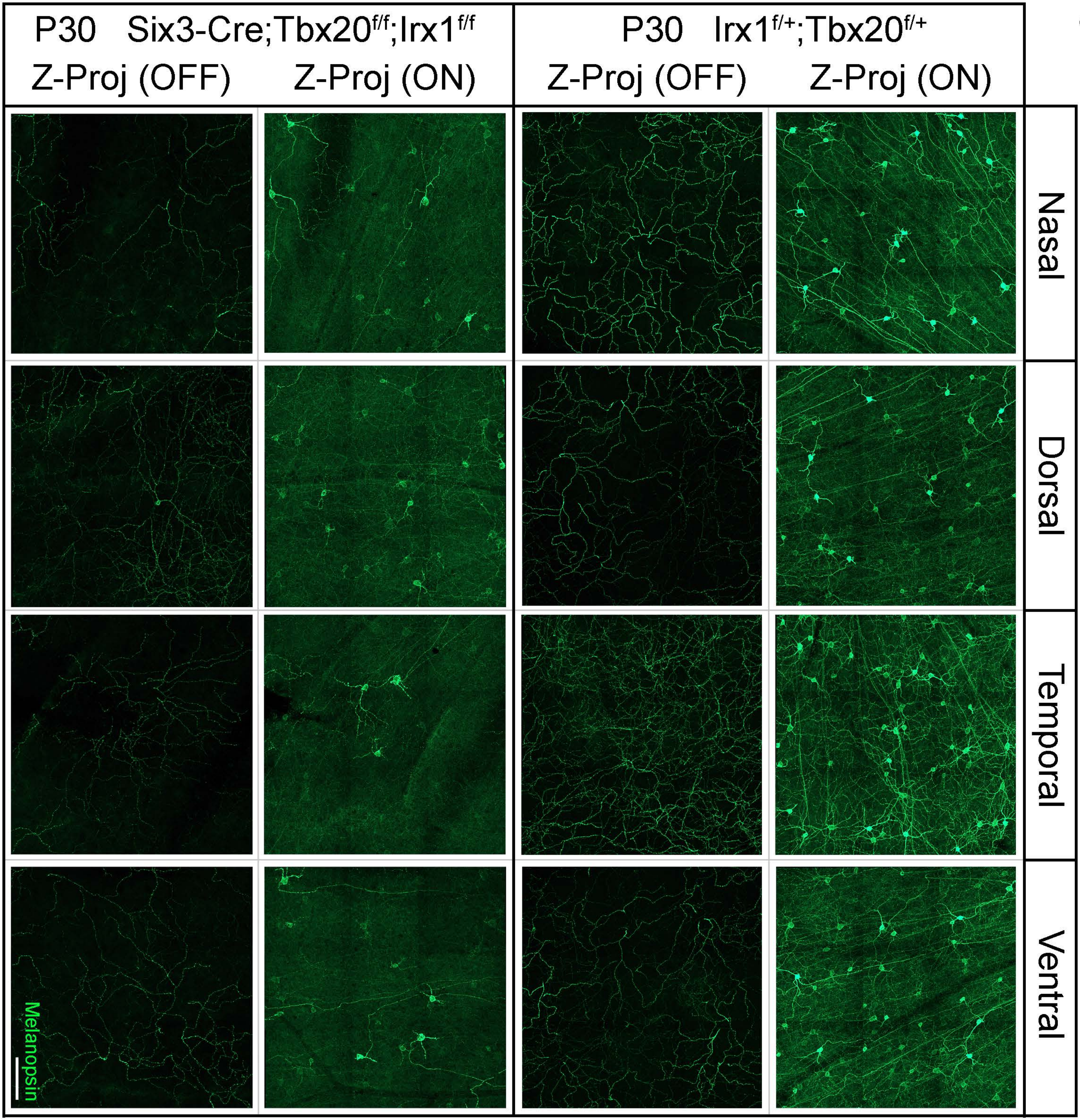

P30 Six3-Cre;Tbx20f/f;Irx1f/f
Z-Proj (OFF)
Z-Proj (ON)
P30 Irx1f/+;Tbx20f/+
Z-Proj (OFF)
Z-Proj (ON)
Nasal
Dorsal
Temporal
Ventral
Melanopsin

Fig. S13

Fig. S14

A

**Irx1**
**mm9 chr13:72,096,064-72,100,936**

**Tbr2-bound element in E14.5 cortex**
**mm9 chr13:72,089,243-72,089,812**

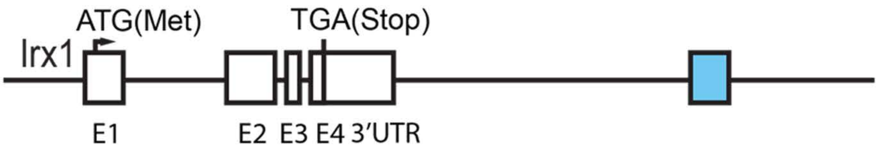


```
>mm9_dna range=chr13:72089243-72089812 5'pad=0 3'pad=0 strand=- repeatMasking=none
AGGACTAACTAGACCCTCTCTGTCTTATTAAAAACCTCTGCTTGCTCCCAGATAATGAAATGAGATTTCTCCTCTCTAGCACTGTCTCCCCTTTTGCCTGCTTTCTA
ATCTCTGGCCACTGTGGTTTTGGGATTACTCCAGCAGGCTGTACAAAGGGAGTCGAAGCCATCTGAAGTGTCACTCCGCGGCTGCCATGAAGCAGCTGCCTTGCCAT
TTTGTTGATGCCCCCAGCTCATGTGTGCAGCAGAAGGTGAAATCAAAGCGGCCCGTCTTCCTCAGTTCTGCACATTATTTTTCACCGCTGTGGCGAATCTCTCCGAG
CTCCGTGAACCAATAAAACAACGGCCCCTGTTTGAGTTGATATATTAATATTTACTGAGGATTTGCAAGGTCGGGGCTAAATGTCAGCAGATTGCTGGGGCAGTCCT
GTGGGGAATAATTAAACACATCTCACAGCAGTTCAGCTTTCTAACATCTTTATTAATCTTTTTTTCCCCTTCCCTCCTGAAAGTGTACATTGAAATATTGCCTCTGC
AACCTCAGAGAAGGCTCAGAATAAGTTTAATAATA
```

JASPAR ID
UN0307.1 EOMES Homo sapiens TBrain-related factors

| Matrix ID | Name | Score | Relative score | Start | End | Strand | Predicted sequence |
|---|---|---|---|---|---|---|---|
| UN0307.1 | UN0307.1.EOMES | 8.771 | 0.865 | 452 | 462 | + | CACAGCAGTTC |
| UN0307.1 | UN0307.1.EOMES | 7.823 | 0.845 | 95 | 105 | - | GAAAGCAGGCA |

B

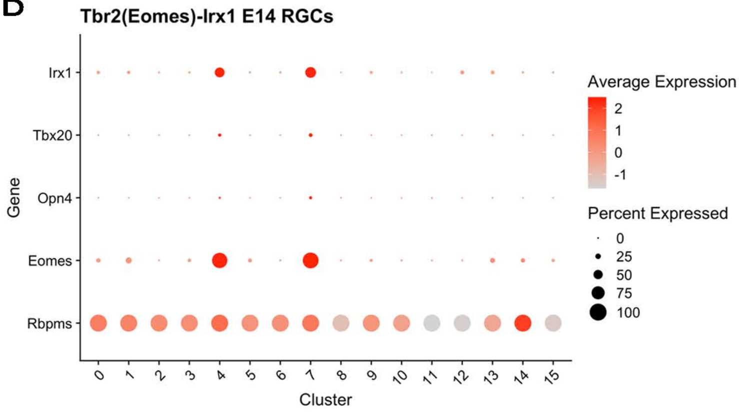


C

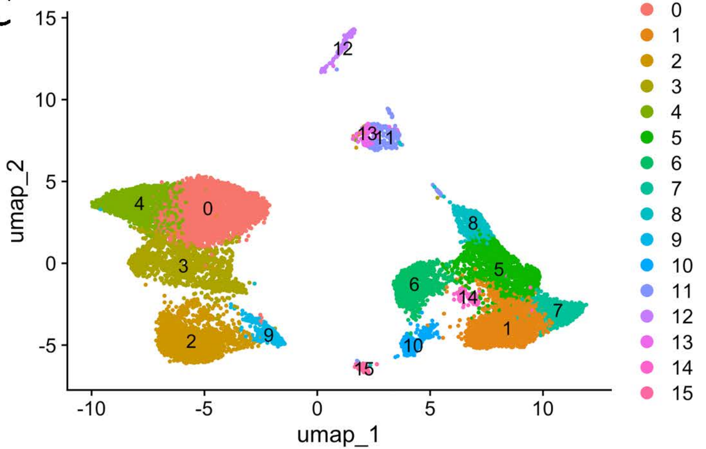
